# Structural and functional basis of the non-canonical human Dicer-tRNA complex

**DOI:** 10.64898/2026.08.12.744379

**Authors:** Arianna Di Fazio, Stephan Hirschi, Federica Battistini, Nuno B. Santos, James Boot, Kamal Ajit, Abdullah Abdullah, Adele Alagia, Modesto Orozco, Monika Gullerova

## Abstract

Human Dicer (hDicer) is a key enzyme in the RNA interference (RNAi) pathway that generates ∼21-22 nt micro-RNA (miRNAs) and small interfering RNAs (siRNAs). We have previously shown that hDicer also generates tRNA-derived small RNAs (tsRNAs), which mediate nuclear gene silencing and regulate hundreds of disease-associated genes. As powerful and evolutionarily conserved cellular regulators, tsRNAs emerged as an important class of small RNAs. Therefore, it is essential to understand their biogenesis. However, the molecular and structural basis of tRNA cleavage by hDicer, as well as the role of chemical modifications such as 5-methylcytosine (m^5^C), in this process, remain unknown. Here, we present the first structural insights into hDicer in complex with tRNA, obtained by cryo-electron microscopy (cryo-EM), selective 2′-hydroxyl acylation analyzed by primer extension (SHAPE) and molecular dynamics (MD) simulations. Our results reveal that tRNAs adopt alternative conformations that are recognized and processed by hDicer. Furthermore, we show that tRNA cleavage by hDicer is facilitated by the m^5^C modification deposited by Nop2/SUN RNA methyltransferase 2 (NSUN2). Collectively, our findings redefine tRNAs as bona fide hDicer substrates and uncover a modification-dependent biogenetic pathway that reshapes the current understanding of the origins and regulation of human small RNAs.

**Graphical Abstract:** 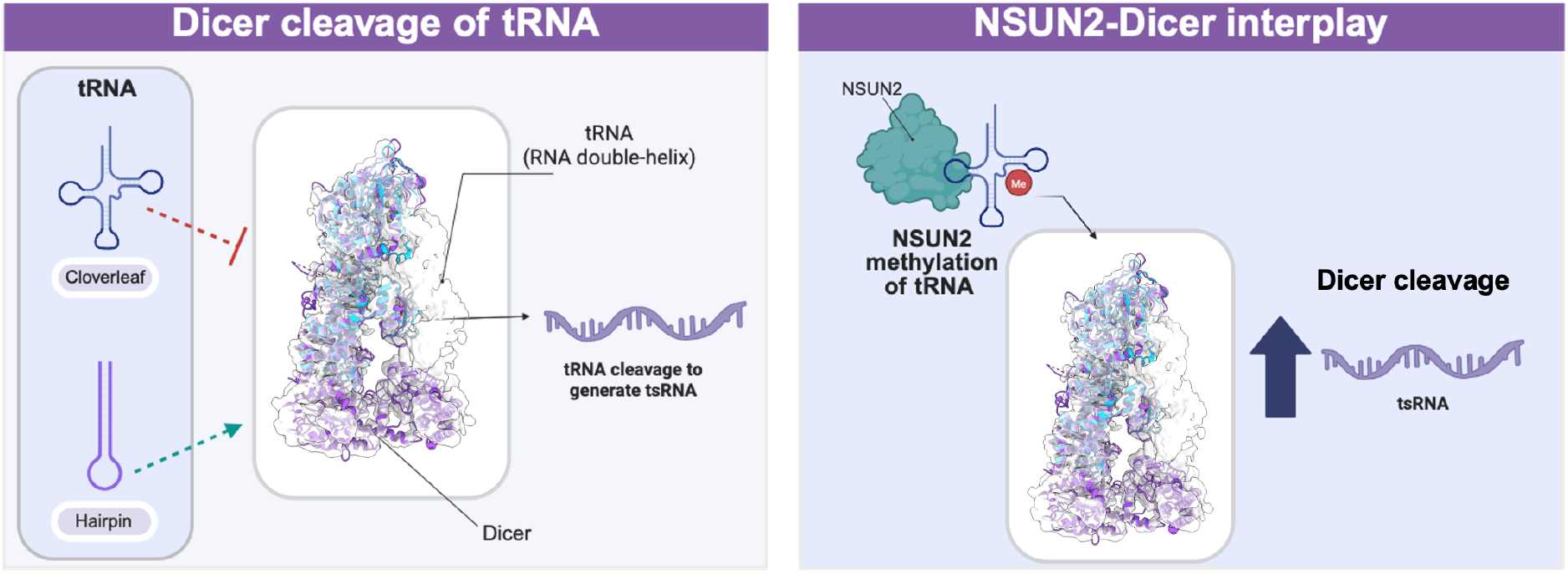

## INTRODUCTION

Small regulatory RNAs act as guide molecules in the RNA interference (RNAi) pathway^1^. Central to this process is the ribonuclease (RNase) III enzyme human Dicer (hDicer)^2,3^, which cleaves RNA substrates to generate miRNA^4^ and siRNA^5^. Structural and biochemical studies have established the mechanism of canonical precursor miRNA (pre-miRNA) processing, which involves domain-specific recognition of RNA motifs and structural elements by the PAZ^6^, platform^7^, helicase^8,9^, double-stranded RNA-binding (dsRBD)^10^ and RNAse domains^11,12^. Previous cryo-electron microscopy (cryo-EM) studies have reconstructed the pre-dicing to dicing transition of hDicer during pre-miRNA processing^13,14^. In the pre-dicing state, the helicase domain engages the terminal loop of the pre-miRNA, restricting access to the processing valley until RNA conformational rearrangements mediated by the dsRBD occur^13^. Subsequent conformational changes in hDicer, including reorientation of the PAZ helix (residues 1014–1029), increase helicase flexibility, enabling transitions between the ‘open’ and ‘closed’ conformations and displacing the dsRBD to facilitate docking of the RNA substrate within the catalytic RNase core^14^.

Accumulating evidence indicates that hDicer also processes a diverse range of non-canonical RNA substrates beyond miRNAs and siRNAs, including vault RNAs (vtRNAs)^15^, small nucleolar RNAs (snoRNAs)^16,17^, damage-responsive transcripts (DARTs)^18^ and R-loops^19^. Transfer RNAs (tRNAs), which are essential for protein synthesis, are likewise cleaved by hDicer to generate tRNA-derived small RNAs (tsRNAs)^20–25^. Deep sequencing studies have identified evolutionarily conserved tsRNAs across vertebrates^24,26^. These molecules have emerged as powerful and conserved regulators of gene expression, cellular stress responses, development and disease, underscoring the importance of understanding their biogenesis. In our previous study, we identified hDicer-derived 3′ tsRNAs carrying CCA tails as *bona fide* regulatory RNAs that mediate a nuclear RNAi mechanism termed nascent RNA silencing^23^. Beyond their regulatory functions, tsRNAs play central roles in the cellular stress response. Their production is strongly induced by oxidative stress, nutrient deprivation, hypoxia, and viral infection, where they promote stress-granule formation, suppress non-essential protein synthesis, and redirect cellular resources toward survival pathways^27–37^. During host–pathogen interactions, tsRNAs can also directly inhibit viral replication or modulate innate immune signalling^38^. Despite their biological importance, the molecular basis of hDicer-mediated tRNA cleavage remains largely unknown.

To investigate the mechanism of hDicer-mediated tsRNA biogenesis, we combined cryo-EM, 2′-hydroxyl acylation analysed by primer extension (SHAPE), and molecular dynamics (MD) simulations. We present the first structural and functional insights into the recognition and processing of non-canonical substrate tRNAs, demonstrating that these molecules adopt alternative conformations that are selectively recognised and cleaved by hDicer.

In cells, tRNAs are extensively decorated with chemical modifications that regulate their maturation, structural integrity and functional fidelity^39,40^. We recently identified Nop2/SUN RNA m^5^C-methyltransferase 2 (NSUN2), which deposits methylcytosine (m^5^C) on tRNAs, as an hDicer-interacting protein^18,41^. To investigate the interplay between NSUN2-mediated m^5^C modification and hDicer-dependent tRNA processing, we show here that m^5^C promotes hDicer-mediated cleavage of tRNAs. Together, our hDicer-tRNA^(m5C)^ structural model provides a molecular mechanism of non-canonical tRNA processing and its regulation by NSUN2.

## MATERIAL AND METHODS

### Protein expression and purification

Proteins were expressed using the Bac-to-Bac expression system (Thermo). pFastBac1 plasmid containing a N-terminal 6xHis-tag, followed by a TEV cleavage site and the cassette for protein of interest was transformed into MAX Efficiency™ DH10Bac Competent Cells *E. coli* (Gibco, 10361012) and colonies screened with the blue/white selection to generate bacmid DNA according to manufacturer’s instructions. The bacmid was isolated from positive colonies by lysing the bacteria with Miniprep kit buffer P1 (Qiagen, 27104) and then isopropanol precipitation, followed by 100% and 70% ethanol washes. Isolated bacmid was then transfected into Sf9 cells grown in static incubator with lipofectamine ExpiFectamine™ Sf Transfection Reagent (Themo Fisher, A38915) by mixing 250 μL of optiMEM to 10 μL expifectamine and 1 μg of bacmid for 0.8 x 10^6^ cells. 2.5 mL of V_0_ baculovirus stock was collected 10 days after transfection and was used to infect 50 mL Sf9 cells and the cells were split every day to 0.5 x10^6^ cells/mL. V_1_ baculovirus stock was collected 48 hours after the cells stopped growing. 1L of low-passage (<30) Sf9 cells were seeded at 0.5 × 10^6^ cells/ml and grown in suspension at 27 °C, 110 rpm. When the cells reached log phase, 1.5–2.5 × 10^6^cells/mL, were infected with 5 mL of V_1_ baculovirus stock. Cells were harvested after 72 hours by centrifugation at 700 g for 15 minutes at 4 °C.

Standard procedure of protein purification of 6xHis-tagged protein with TEV cleavage site was as follows, with all steps carried out at 4 °C. Cell pellets were lysed with 6 volumes of lysis buffer (20 mM Tris–HCl, pH 8, 500 mM NaCl, 10% glycerol, 0.5% Triton X, 1 mM TCEP, 1x Complete protease inhibitor EDTA-free, 1x PMSF, and 1x leupeptin) and 2500 U of universal nuclease (Pierce), with rotation for 30 min. The lysate was then sonicated five times for 30 s on/off at 15 μm, spun at 13 000 *g* for 45 min, and the supernatant filtered with a 0.2-μm filter (Corning, 430049). Cleared lysate was then loaded onto Ni-NTA agarose resin (QIAGEN, 30210) in a gravity flow column pre-equilibrated with 5 volumes of wash buffer and washed with 20 volumes of wash buffer (20 mM Tris–HCl, pH 8, 500 mM NaCl, 5% glycerol, 1 mM TCEP, 1x Complete protease inhibitor EDTA-free, 1x PMSF, 1x leupeptin, and 10 mM imidazole). Proteins were eluted with elution buffer (20 mM Tris–HCl, pH 8, 500 mM NaCl, 5% glycerol, 1 mM TCEP, 1x Complete protease inhibitor EDTA-free, 1x PMSF, 1x leupeptin, and 500 mM imidazole). Fractions were run on SDS–PAGE and stained with Coomassie blue. hDicer-containing fractions were dialysed against buffer (50 mM Tris–HCl, pH 8, 300 mM NaCl, 10% glycerol) overnight and the tag was cleaved with TEV Protease (NEB, P8112S) or ProTEV Plus (Promega, V6101) during dialysis. Reverse nickel purification was carried out to remove uncleaved hDicer by incubating the sample for 10 minutes with equilibrated Ni-NTA agarose resin; the flow-through was concentrated with a 50 kDa or 100 kDa cut-off concentrator (Vivaspin, 28-9323-63) and subjected to size exclusion chromatography using a Superdex 200 Increase 3.2/300 column mounted on an ÄKTA Pure (Cytiva). Active fractions were pooled, concentrated and stored in 50 mM Tris–HCl (pH 8), 150 mM NaCl, 5 mM MgCl_2_, 1 mM DTT, 20% glycerol, at −80°C.

### *In vitro* NSUN2 methylation reaction and Dicer-RNA immunoprecipitation (RIP)

10 μg of tRNA (400 pmol) obtained from *in vitro* transcription was incubated with 1200 pmol of recombinant NSUN2 protein purified from Sf9 cells, at 37 °C for 2 hours in 50 mM Tris-HCl (pH 8), 300 mM NaCl, 5 mM MgCl_2_, 5% glycerol, 40 U of RNAse (ribolock), 1 mM DTT, 1.5 mM SAM. 200 pmol of purified hDicer^DEDE^ were added in the reaction and incubated on ice for 60 minutes. Reaction was pre-cleared with 50 μL of Dynabeads Protein G (Life Technologies, 10004D) with rotation at 4 °C for 60 minutes. After pre-clearing, reaction was incubated with 7.5 μg of anti-Dicer antibody (Abcam, 14601 and Supplementary Table 2) with rotation (18 rpm) at 4 °C for 2 hours. 200 μL of pre-equilibrated beads were allowed to bind for 1.5 hours at 4 °C. Beads were then washed 5 times with wash buffer (50 mM Tris-HCl pH 8, 150 mM NaCl, 1 mM MgCl_2_, 0.05% NP40, 1mM DTT, 1x protease inhibitor and 200 U/mL RNase inhibitor) and were incubated with proteinase K for 30 minutes at 37 °C and shaking at 700 rpm. RNA was incubated with Trizol by shaking at 65 °C for 5 minutes, followed by standard Trizol extraction as described earlier.

### Oxford Nanopore Direct RNA sequencing

### Library preparation

RNA samples were diluted in DEPC-treated water and quantified with Qubit using the RNA HS Assay kit (Invitrogen, Q32852). 50 ng of RNA were heated at 95 °C for 3 minutes and immediately placed on ice to disrupt secondary structures. RNA was polyadenylated with 15 U of *E. coli* poly(A) polymerase (NEB, M0276S) and 1 mM ATP in 1x E. coli poly(A) polymerase buffer for 60 seconds at 37 °C. Reaction was quenched by adding EDTA to a final concentration of 10 mM. The reaction was then mixed with 2.25 volumes of Agencourt RNAClean XP beads (Beckman Coulter, A63987), resuspended and incubated at room temperature for 5 minutes. Beads were then washed very gently with 300 μL of ice-cold 80% ethanol once. RNA was eluted in 26 μL of DEPC-treated water for 10 minutes by incubating at room temperature with frequent vortexing. The following steps were carried out with the Direct RNA sequencing kit (SQK-RNA004). Reverse transcription adapters splint ligation was carried out with 4M U of T4 DNA Ligase (NEB, M0202M) with 1 μL of RT adapters (RTA) and 1.25 μL of RNasin plus RNase inhibitor (Promega, N2615) in NEBNext Quick ligation reaction Buffer (NEB, B6058S), in a total reaction of 35 μL, and incubated at 23 °C for 10 minutes. Reverse transcription was carried out by adding 200 μM dNTPs and 400 U of Induro RT (NEB, M0681S) in 1x Induro RT reaction buffer added directly in the previous reaction and was incubated at 60 °C for 30 minutes and then 70 °C for 10 minutes. RNA was cleaned with 2 volumes of RNA XP beads as described earlier and resuspended in 33 μL of DEPC-treated water. 6 μL of sequencing adapters (RLA) were ligated with 6M U of T4 DNA Ligase in NEBNext Quick Ligation Reaction Buffer and incubated at 23 °C for 20 minutes. RNA was isolated with 2 volumes of RNA XP beads, washed with Wash Buffer (WSB) and eluted in 20 μL of RNA Elution Buffer (REB). Eluted RNA library was mixed with Sequencing buffer (SB) and Library Solution (LIS) according to manufacturer’s instruction. Finally, RNA samples were loaded on RNA flow cell (ONT, FLO-MIN004RA) primed as per manufacturer’s instructions and mounted on minION (ONT).

### Data processing

Base call of the Nanopore data was carried out using DORADO (https://github.com/nanoporetech/dorado) with the latest ‘Super High Accuracy’ base calling model released by ONT. RNA modification models for Ψ, m5C, m6A, and Inosine were used in the base calling step. The DORADO Aligner was used to align the base-called reads to the FASTA sequence of mature tRNA-Pro-TGG with the parameters -ax ont -k 14. Modified residues were then annotated using the MODKIT pileup command, with the modification threshold set at 0.95 for all samples. The bedMethyl table produced from this command contains the percentage of m5C-modified cytosines across the mature tRNA-Pro-TGG sequence. A custom Python script was used to plot the bedMethyl table, showing the percentage of m^5^C across the tRNA sequence.

### Selective 2′-Hydroxyl Acylation Analyzed by Primer Extension RNA probing

#### Sample and library preparation

tRNA-Alanine-AGC-2-1 and tRNA-Pro-TGG-3-5 were *in vitro* transcribed and resuspended to 125 ng/μL (∼5 μM) in DEPC water. 20 μL of the tRNAs were annealed by heating the samples at 95 °C for 2 minutes then mixed with 10 μL of 3.3 X annealing buffer (333 mM Hepes pH 8, 333 mM NaCl, 33 mM MgCl_2_) and incubated at 37 °C for 20 minutes to allow folding. 0.8 μM of folded tRNA was incubated with 7 molar excess hDicer or equal volume of protein storage buffer on ice for 30 minutes to allow binding. The binding reactions were reacted with SHAPE reagents 5NIA (27.5 mM) or DMS (1% v/v) for 15 min and 5 min respectively at 37 °C, or equal volume of DMSO as negative control for 15 min at 37 °C. Reactions were quenched with 120 mM DTT (5NIA) or 133 mM DTT (DMS/DMSO). Reacted RNA was extracted with 6 volumes of Trizol. To purify the RNA, 0.2 volumes of chloroform were added to the samples in TRIzol, followed by vortexing for 3 min. Phases were separated by centrifuging 15 min at 15 000 g and 4°C, and the aqueous phase was collected to a new tube. One volume of 100% EtOH was added and samples were purified using Direct-zol RNA Microprep Kit (Zymo Research, R2060), according to manufacturer’s instructions. After elution, samples were incubated at 95 °C for 3min, and subsequently on ice for 3 min. PolyA tailing was performed using E. coli Poly(A) Polymerase (NEB, M0276), followed by clean-up using RNA Clean & Concentrator-5 (Zymo Research, R1013), both according to manufacturer’s instructions. RNA samples were diluted in 6.37 µl nuclease-free water, added to 2 µl of 10mM dNTP mix (NEB, N0447S) and 0.68 µl of 100 µM RT primer (CTACACGACGCTCTTCCGATCTNNNNNNNNNNTTTTTTTTTTTTTTTTTTTTTGG). Samples were incubated at 65 °C for 5 min, followed by another incubation on ice for 2 min, and then diluted in reaction buffer (final concentration 50mM Tris-HCl (pH 8.0), 75mM KCl, 6mM MnCl2, 10mM DTT, 10U SUPERase·In™ RNase Inhibitor (ThermoFisher, AM2694) and 1M betaine). After an incubation at 25 °C for 2 min, first-strand synthesis was performed by adding 200 U of SuperScript II (Invitrogen, 18064022), with following program: 25 °C, 10 min; 42 °C, 90 min; 10 x 50 °C, 2 min, 42 °C, 2min; 70 °C, 15 min. Samples were purified with 2 volumes of AMPure XP beads (Beckman Coulter, A63880), following manufacturer’s instructions. Subsequently, samples were prepared using xGen™ RNA Library Prep Kit (IDT, 10009814), according to manufacturer’s instructions (starting from ExoI treatment), until the final indexing PCR, where NEBNext® Multiplex Oligos for Illumina® (NEB, E6609S) were used. Finally, sequencing was performed in an Illumina NovaSeq X Plus.

#### Data processing

Raw sequencing reads (R1 only) from all lanes and experimental runs were first concatenated. A pre-processing pipeline was then applied, which involved trimming adapter sequences and poly-G tails from dark cycles (cutadapt v4.2), extracting unique molecular identifiers (UMIs) (umi_tools v1.1.4), hard-clipping the adaptase sequence (seqtk v 1.4), and reverse-complementing the reads (fastx_toolkit_0.0.14). After pre-processing, read quality was assessed using FastQC v0.11.8 and MultiQC v1.25.1. The processed reads were subsequently aligned to the corresponding tRNA sequence for each sample, using Bowtie2 v2.5.1 in very-sensitive-local mode with parameters optimized for SHAPE/DMS-MaP analysis (--mp 3,1 --rdg 5,1 --rfg 5,1 --dpad 30). Following alignment, PCR duplicates were removed based on their UMIs using umi_tools. The reactivity profiles were generated using the RNAFramework pipeline. First, the count module was used to calculate per-nucleotide reverse transcription (RT) stop/mutation rates and read coverage. Reactivity scores were then calculated using the norm module by taking the difference between the normalized values for the modified and control samples within each tRNA transcript sample. Normalisation was performed by dividing the natural logarithm of the raw count at each position by the natural logarithm of the average count across the transcript. The final reactivity score was set to the maximum of zero or the calculated difference between the treated and untreated samples. These reactivity scores were converted to a wiggle format using the wiggle module and used as pseudo-energy constraints to predict the RNA secondary structure with the fold module, which utilizes the ViennaRNA method. Finally, to generate a single, robust structural model for downstream analysis, reactivity profiles from all replicates were merged using the combine module before the final folding step. Coverage plots were generated by collecting coverage using the rf-rctools view module of RNAFramework, importing coverage text files into R-4.5.0 and plotting using ggplot2. SHAPE plots were generated using a custom script (https://github.com/Mearsh/shapeplotr). Delta-SHAPE plots for the comparison of Pro and ProDic samples were generated by loading reactivities for each relevant sample into R-4.5.0, calculating mean and standard deviation for replicate reactivities and then calculating the difference between mean and standard deviation of reactivities between tRNAs of interest, for each treatment group (DMS and 5NIA), plots were generated using ggplot2.

### *In vitro* reconstitution of the Dicer-tRNA^-m5C^ complex

tRNA-Proline-TGG 3-5 was *in vitro* transcribed and folded by slow cooling. To allow methylation, 6 μM of tRNA-Pro was incubated with 3 μM of purified NSUN2-WT for 1 hour at 37 °C in 50 mM Tris-HCl (pH 8), 150 mM NaCl, 10 mM MgCl, 0.5 mM TCEP, 5% glycerol, 1.5 mM SAM. Following, 4.5 μM of purified hDicer^DEDE^ and 3 mM CaCl_2_ were added to the reaction and incubated on ice for 30 min to allow Dicer-tRNA binding. The assembled complex was then separated by size exclusion chromatography on a Superdex Increase 3.2/300 column (Cytiva) in 50 mM Tris-HCL, 100 mM NaCl, 0.5 mM TCEP, 2 mM CaCl_2_. Fractions containing the complex (B9-B1, Supplementary Fig. S3B) were collected, pooled and concentrated to 1.2 mg/mL with a 100 KDa cutoff spin column (Proteintech,88503).

### Structure determination using cryo-EM

#### Cryo-EM sample preparation and data collection

3 μl of purified hDicer-tRNA-NSUN2 complex at 1.2 mg/ml were applied to C-Flat 1.2/1.3 grids, that had been glow-discharged for 60 s at 15 mA. Grids were blotted for 4-6 s and immediately vitrified by plunging into liquid ethane using an FEI Vitrobot Mark IV (Thermo Fisher) at 4 °C and 100% humidity. Data were collected on an FEI Titan Krios operated at 300 kV and equipped with a Gatan BioQuantum energy filter (20 eV) and a K3 direct electron detector using EPU (Thermo Fisher). 48,488 micrograph movies were collected from two separate grids prepared with the same sample. Images were acquired at a magnification of 58,149x, corresponding to a pixel size of 0.83 Å/px, using a 100 μm objective aperture and a defocus range of -0.5 to -2.5 μm with an accumulated dose of 42.8 e^-^/Å^2^ over 40 frames.

#### Cryo-EM data processing

The cryo-EM data processing workflow is illustrated in Supplementary Figure S4. Patch-based motion correction of movies was performed on-the-fly using SIMPLE 3.0^42^ followed by patch CTF estimation and manual micrograph curation in cryoSPARC v4.7.1^43^ during which 45,305 micrographs were selected. An initial set of particles was picked from a subset of micrographs in cryoSPARC using the blob picker to generate templates for automated picking using the template picker. In parallel, micrographs were also picked using TOPAZ^44^ with the previous templates as input. The two resulting particle sets were then merged and, after removal of duplicates, yielded a total of 19,386,122 unique particles. After multiple rounds of stringent 2D classification, to remove low-quality particles and other non-protein contaminants, a final stack of 514,000 particles was obtained. Notably, at this stage, neither heterogeneous refinement or 3D classification after local refinement in cryoSPARC, nor 3D classification with or without Blush regularisation in Relion 5.0.0^45,46^ was able to resolve the remaining particles into different classes. This, together with the blurry tRNA region visible in 2D classes (Supplementary Fig. S4) suggested the presence of continuous heterogeneity, which was attempted to be resolved using 3D variability analysis as well as 3D flex refinement in cryoSPARC (Supplementary Videos 1 and 2). However, no further classification of discrete states was possible. The final stack of particles was exported to Relion for Bayesian particle polishing, before a final round of 2D classification and non-uniform refinement in cryoSPARC with 485,482 particles yielded a 3.33 Å consensus map of the full particle. To further improve the different regions of the map, local refinements with masks focusing on 1) hDicer plus tRNA without the helicase domain (helicase-less mask) and 2) the PAZ and Platform domains plus tRNA (PAZ-tRNA mask) were performed. This yielded more detailed local maps with resolutions of 3.32 Å using the helicase-less mask and 3.47 Å using the PAZ-tRNA mask. To aid manual model building in Coot^47^, maps were sharpened using DeepEMhancer^48^ and a composite map was generated from consensus and local refinement maps in ChimeraX^49^. Local resolution was estimated using MonoRes^50^ in cryoSPARC at an FSC threshold of 0.143. Particle Euler angles, local resolution and Fourier shell correlation (FSC) plots are presented in (Supplementary Fig. S5). ChimeraX was used to render and illustrate all maps and models^49^.

#### Model building and refinement

To facilitate model building, the cryo-EM structure of human hDicer in a pre-dicing state (PDB ID: 5ZAL)^13^ was used as a starting point. It was docked into the consensus map, and the following mutations were introduced: D1320A; E1444A; D1709A; E1813A. The final protein model was obtained by multiple iterations of manual model building in Coot and real-space refinement in PHENIX^51^. MolProbity^52^ and PHENIX were used for structure validation, and all statistics were compiled in Table 1 together with information about the cryo-EM data acquisition and processing. The final protein model was deposited in the Protein Data Bank (PDB ID: 9TE0) and cryo-EM maps in the Electron Microscopy Data Bank (EMDB ID: EMD-55806, EMD-55807, EMD-55808, EMD-55809).

**Table 1.** Cryo-EM data collection, refinement and validation statistics.

| Dicer-tRNA complex<br>(EMD-55807, PDB 9TE0) |  |
| --- | --- |
| **Data collection** |  |
| Microscope | Titan Krios |
| Camera | Gatan Quantum-K3 |
| Voltage (kV) | 300 |
| Magnification | 58,149x |
| Defocus range (μm) | -0.5 to -2.5 |
| Movies recorded | 48,488 |
| Frames per movie | 40 |
| Total dose per movie (e<sup>-</sup>) | 42.8 |
| Pixel size (Å) | 0.83 |
| **EM data processing** |  |
| Initial particles picked | 19,386,122 |
| Particles used for | 485,482 |
| Box size (Å) | 299 Å |
| Symmetry imposed | C1 |
| Consensus map | 3.3 Å |
| Map resolution range | 2.1 – 11.9 |
| Map sharpening method | DeepEMhancer<sup>7</sup> |
| **Model refinement and validation** |  |
| Initial model used | PDB: 5ZAL |
| Model composition |  |
| Chains | 1 |
| Protein residues | 769-883, 891-950, 971-1072, 1292-1378,<br>1551-1583, 1663-1781, 1803-1910<br>(624 total) |
| Ligands | 0 |
| Waters | 0 |
| RMSD |  |
| Bond length (Å) | 0.003 |
| Angles (°) | 0.622 |
| Molprobity score <sup>c</sup> | 2.17 |
| Clash score | 16.33 |
| Rotamer outliers (%) | 0 |
| Ramachandran |  |
| Favoured | 92.85 |
| Allowed | 7.15 |
| Outliers | 0.00 |
| Ramachandran Z-score | -1.52 |
| CaBLAM outliers (%) | 3.73 |
| EM-Ringer score <sup>d</sup> | 0.69 |
| Map CC (peak) <sup>e</sup> | 0.31 |
| Map CC (mask) <sup>e</sup> | 0.53 |

<sup>a</sup> Resolution determined by FSC with cut-off of 0.143. <sup>b</sup> Resolution range defined by local resolution values at atom positions in Chimera<sup>49</sup> <sup>c</sup> Model statistics were calculated using MolProbity<sup>52</sup> <sup>d</sup> Calculated based on local fit of side chains to map according to Barad *et al.*<sup>62</sup> <sup>e</sup> Real-space correlation coefficients of model-to-map fit as described in Afonine *et al.*<sup>63</sup>

### Molecular Dynamics

All molecular dynamics (MD) simulations were performed using AMBER 24. The starting structures were based on the cryo-EM map of Dicer built on PDB:5ZAL without the helicase domain, and model tRNA-Pro obtained with trRosettaRNA^53^ based on SHAPE. The protein was parameterized with the ff19SB force field^54^, while RNA was described with the OL3 force field^55^. Water molecules were modelled with the TIP3P model^56^. Ion parameters were taken from Joung et al.^57^. Each system was solvated in an octahedral TIP3P water box with a 10 Å buffer and neutralized with K+ counterions. Additional K+ and Cl− ions were added to mimic a physiological salt concentration of ∼0.15 M. The systems underwent standard optimization procedures, consisting of a five-step energy minimization (first restraining solute atoms with a 25 → 5 kcal·mol−1·Å−2 force constant, followed by unrestrained minimization). This was followed by a gradual thermalization to 298 K. Subsequently, simulations were carried out under constant pressure and temperature (NPT ensemble, 1 atm, 298 K). Finally, production MD simulations of 1 μs were performed for each system under NPT conditions (1 atm, 298 K) without positional restraints.

### RNA *in vitro* transcription

DNA templates for *in vitro* transcription were purchased as ssDNA ultramers (4 nmoles) from IDT, containing the T7 polymerase promoter sequence (TAATACGACTCACTATA<u>GGG</u>), -GGG were included to increase transcription yield. All IVT tRNA sequences were designed with -CCA triplet at the 3′ end (tRNA-Proline: TAATACGACTCACTATAGGGGGCTCGTTGGTCTAGGGGTATGATTCTCGCTTTGGGTGCGAGAGGTCCCGGGTTCAAATCCCGGACGAGCCCCCA; tRNA-Alanine: TAATACGACTCACTATAGGGGGGGGTGTAGCTCAGTGGTAGAGCGCGTGCTTAGCATGCACGAGGCCCCGGGTTCAATCCCCGGCACCTCCACCA; pre-let7a: TAATACGACTCACTATAGGGTGGGATGAGGTAGTAGGTTGTATAGTTTTAGGGTCACACCCACCACTGGGAGATAACTATACAATCTACTGTCTTTCCTA). Equimolar amounts of antisense and sense DNA strands were annealed in DNA annealing buffer (10 mM Tris–HCl pH 7.6, 50 mM NaCl and 1 mM EDTA pH 8), by incubating the DNA at 95°C for 3 min, followed by slowly cooling down to room temperature. RNA IVT was carried out with HiScribe T7 High Yield RNA Synthesis Kit (NEB, E2040S) with 2 μg of dsDNA template, NTPs (10 mM each), 2 μL of T7 polymerase mix in 1x rection buffer in a total volume of 20 μL. Reaction was incubated at 37°C for 16 hours; DNA template was then degraded with Turbo DNAse (4 U, Invitrogen, AM2238) for 1 h at 37°C. RNA was isolated from each reaction with 400 μL of TRIzol LS reagent (Invitrogen) and 200 μL of 1-bromo-3-chloropropane (Sigma, B9673-200ML), and then again with 400 μL of 1-bromo-3-chloropropane. RNA was then precipitated in 1 volume isopropanol and 0.75 M ammonium acetate overnight at -20 °C and washed with 100% ethanol, followed by a 70% ethanol wash. RNA pellets were air dried and resuspended in DEPC-water. For all additional oligonucleotides see Supplementary Table 1.

### Electrophoretic mobility shift assay

*In vitro* transcribed tRNA-Pro, tRNA-Ala or pre-let7a (1.6 μM) were incubated with hDicer^DEDE^ (0, 0.25, 0.5, 1, 1.5 μM) in binding buffer (30 mM Tris-HCl pH 7.6, 25 mM NaCl, 2 mM MgCl_2_, 1 mM DTT, 1 mM EDTA) for 45 minutes on ice. RNA-protein complexes were mixed with 1x Native agarose gel loading dye (Invitrogen) prior to separation on polyacrylamide TBE-PAGE gel (6%) in 1xTBE, run at 120V at 4°C. Bands were visualized with a UV transillumination in a Gel Doc™ XR+ imager after staining with SYBR™ Gold.

### Proximity Ligation assay

Cells were fixed and permeabilised at room temperature for 10 min each with 4% paraformaldehyde in PBS (Alfa Aesar, J61899) and 0.2% Triton X-100 (Sigma, X100-500ML) in PBS. Duolink Proximity Ligation Assay (Sigma Aldrich, DUO92101-1KT) was carried out according to manufacturer’s protocol. Coverslips were mounted using Duolink In Situ Mounting Medium with DAPI (Sigma-Aldrich, DUO82040) and stored protected from light at 4°C. Slides were imaged using Olympus Fluoview Spectral FV1200 confocal microscope.

### Foci quantification

Images were imported in FiJi software, z-stacks were generated by the z-project function and converted to 8-bit format. Threshold was set as default, and cell nuclei were identified with the ‘find particles’ function with size settings 100-∞. Foci per nucleus were identified with the ‘find maxima’ function with a cutoff of 40, after having selected only the area occupied by a single cell nucleus. Cells at the edges of the frame were excluded.

### *In vivo* FLAG-Dicer RNA Immunoprecipitation (RIP)

Approximately 10x10^6^ HEK293T cells in a 15-cm dish were reversed transfected with 35 μL of lipofectamine 3000 and 25 μg of a plasmid expressing FLAG-empty vector or FLAG-Dicer. 48 hours after transfection, cells were incubated with 10 μM NSUN2 inhibitor MY-1B^58^ (Cambridge Bioscience Ltd) or equivalent volume of DMSO as vehicle control for 4 hours. Cells were collected, washed with ice-cold PBS twice and UV-crosslinked at 254 nm at 150 mJ/cm^2^. Cells were lysed with buffer (50 mM Tris-HCl pH 7.5, 150 mM NaCl, 15 mM MgCl_2_, 0.5% Triton X-100, 5% glycerol, 1x protease inhibitor, 80 U Ribolock, 8 U Turbo DNase, for 30 min or a rotating wheel at 4°C, sonicated twice at 10 μm for 10 seconds, and centrifuged at 17 000 x g for 10 minutes. The supernatant was incubated with 100 μL of equilibrated anti-FLAG M2 magnetic beads (Sigma) for 1 hour at 4°C. Beads were then washed twice in high salt buffer (50 mM Tris-HCl pH 7.5, 500 mM NaCl, 15 mM MgCl_2_, 0.05% Triton X-100, 5% glycerol, 1x protease inhibitor), twice in low salt buffer (50 mM Tris-HCl pH 7.5, 150 mM NaCl, 1 mM MgCl_2_, 0.01% Triton X-100, 1x proteases inhibitor) and twice in proteinase K buffer (100 mM Tris-HCl pH 7.5, 50 mM NaCl, 10 mM EDTA). Beads were incubated for 1 hour at 37°C with 0.5 U of thermolabile proteinase K (NEB) at 1000 rpm, and then RNA was extracted with Trizol reagent, following standard phenol/chloroform extraction. Extracted RNA was used as template for reverse transcription, and qRT-PCR. ERCC RNA was used as spike-in internal control.

### Surface Plasmon Resonance

#### Chemical protein biotin-labelling

Recombinant Dicer-DEDE purified by size exclusion chromatography (SEC) was buffer exchanged with 5 consecutive washes with ice-cold PBS in a mini concentrator (Thermo, 88504) with 50 KDa cutoff by spinning at 15 000 g for 5 min. Protein in PBS was then incubated with 20-fold molar excess of EZ-Link NHS-PEG4-Biotin reagent (Life Technologies Ltd, A39259) for 1 hour and 30 minutes on ice. Protein-reagent solution was then loaded on Slide-A-Lyzer™ MINI Dialysis Devices, 3.5K MWCO (Fisher Scientific UK Ltd, 10784274) overnight to equilibrate with SPR running buffer and to eliminate unreacted biotin.

Streptavidin was immobilised on CM5 chip (Cytiva, 29104988) by standard amine-coupling until ∼13 000 RU were reached. Biotin-labelled hDicer^DEDE^ (0.2 mg/mL) was injected at 10 μL/min and captured on the chip via streptavidin affinity in flow cell 2 (FC2) until ∼1500 RU were reached. RNA analytes (IVT tRNA-Pro and pre-let7) were folded in vitro, then diluted in serial 2-fold dilutions (10-0.156 μM) and injected at 10 μL/minutes with 30 s association and 60 s dissociation phases. NSUN2 analytes alone or in combination with SAM and tRNA reactions were pre-incubated at 37 °C for 30 minutes, 2-fold diluted (2.5-0.156 μM) and injected at 10 μL/min with 40 s association and 300 s dissociation phases. For the NSUN2 analytes, after each cycle, regeneration buffers (3M NaCl and 0.3% SDS) were injected at 30 μL/min for 120 s. Measurements were carried out in Biacore T200 SPR system (Cytiva) at 37 °C with running buffer (20 mM Hepes pH 8, 150 mM NaCl, 1 mM DTT, 10 mM MgCl_2_ and 10% glycerol). Reference flow cell (FC1) baseline signal was subtracted to binding signals and subsequentially corrected by subtracting values of blank buffer, injected at the beginning and end of each analyte cycle. Equilibrium analysis was performed by plotting RU at equilibrium versus concentration of the analyte, curves were fitted usign GraphPad.

#### NSUN2-hDicer *in vitro* cleavage assay

Approximately 7.5 pmol (180 ng) of IVT tRNA or synthetic tRNA^m5C^ were heated at 95 °C for 2 minutes, cooled down for 10 minutes at room temperature and then incubated with an equimolar concentration of NSUN2 in reaction buffer (50 mM Tris-HCl pH 7.5, 150 mM NaCl, 10 mM MgCl_2_, 1 mM DTT, 10% glycerol) supplemented with 1.5 mM SAM and 40 U of RNase (Ribolock) and incubated at 37 °C for 60 minutes. Equimolar concentration of hDicer-WT or hDicer^DEDE^ were added and reaction incubated for 60 minutes. Reactions were stopped by mixing with denaturing 1x TBE-formamide dye and stored at -80 °C for Northern blotting. For qRT-PCR, samples were prepared in the same way, but 25 ng of unrelated RNA were used as internal control for each reaction. After hDicer cleavage, samples were treated with Thermolabile Proteinase K (NEB, P8111S) for 30 minutes at 37 °C, followed by heat inactivation at 55°C for 10 minutes and used directly for reverse transcription with primer R1(TGGGGGCTCGTCCGG). cDNA was diluted 1:1000 and then amplified with primers R1 and F1 (GGCTCGTTGGTCTAGGGGTATG) for qPCR. qPCR data were analysed using raw Ct values.

#### Northern blot

Samples were separated on a 10% denaturing (50% g/v urea) polyacrylamide gel in 1xTBE. RNA was transferred onto Hybond N+ positive charged nylon membranes (GE Healthcare, 10506125) by semidry electroblotting in 0.5x TBE buffer at 5 V for 60 minutes followed by UV-crosslinking at 120 mJ/cm2 in a UV crosslinker (Carl Roth) and pre-hybridized in ULTRAhyb-Oligo buffer (Thermo Fisher Scientific, AM8663) at 42 °C for at least 30 min. DNA probes (Sigma) (10 μM) were incubated with 20 U of T4 polynucleotide kinase (NEB,M0201L) or T4 polynucleotide kinase (3′ phosphate minus) (NEB, M0236L) in 1x reaction buffer with 2 μL of 32g-ATP(EasyTides. Adenosine 5′-triphosphate, [gamma-32P]) supplied in 50mM Tricine (pH 7.6) at 6000Ci(222TBq)/mmo (Revvity, BLU502Z250UC) and incubated at 37 °C for at least 60 minutes. Probes were then purified with a G-25 Sephadex column (Cytiva, 27-5325-01) by spinning at 700 g for 2.5 minutes. Radio-labelled probes were hybridised onto the membrane at 42 °C followed by 2 washes (15 minutes each) with 0.1x SCC washing buffer. The blot was exposed to Hyperfilm MP (VWR, 28-9068-45).

#### Statistical analyses

Statistical analyses were performed in GraphPad Prism. Shapiro-Wilk test was performed to assess normal distribution. If the data met the criteria for a normal distribution, statistical analysis was performed using unpaired t-test (two-tailed) or one-way ANOVA with Tukey’s multiple comparison test. For data that did not follow a normal distribution, non-parametric tests including Kruskal-Wallis test with Dunn’s multiple comparisons test were performed to test for significant difference (P-value < 0.05 is considered as significant).

## RESULTS

### hDicer binds tRNA comparably to pre-let7

Endogenous hDicer binds non-canonical substrates, including tRNAs^17,59^. To compare the binding of tRNAs with that of canonical substrate, we measured the affinity of hDicer for the Proline tRNA isotype TGG 3-5 (tRNA-Pro), previously shown to be cleaved by hDicer into functional tsRNAs^23^, and for pre-let7a, a well-characterized canonical miRNA precursor. To determine whether hDicer binds only processable substrates, we also selected Alanine tRNA isotype AGC 2-1 (tRNA-Ala), which is not processed by hDicer *in vivo*^23^. All RNA substrates were synthesized by T7-*in vitro* transcription (Supplementary Fig. S1A and S1B). We purified and chemically biotinylated the catalytically inactive hDicer^DEDE^ mutant carrying the substitutions D1320A, E1444A, D1709A and E1813A ^14,17,60,61^ (Supplementary Fig. S1C and S1D) and immobilized the protein on a streptavidin-coated surface plasmon resonance (SPR) sensor chip. SPR measurements using serial dilutions of tRNA-Pro, tRNA-Ala and pre-let7a revealed comparable binding affinities for hDicer^DEDE^ , with dissociation constants ranging from 8 to 11 μM (Fig. 1A). Electrophoretic mobility shift assays (EMSAs) of hDicer^DEDE^ complexes with tRNA-Pro, tRNA-Ala and pre-let7a further confirmed binding of hDicer to all three RNA substrates *in vitro* (Fig. 1B). To validate these interactions in cells, we overexpressed FLAG-hDicer in HEK293T cells and performed RNA immunoprecipitation (RIP). Consistent with the *in vitro* results, hDicer associated with all three RNA substrates, including tRNA-Pro, tRNA-Ala and pre-let7a (Fig. 1C).

**Figure 1.**
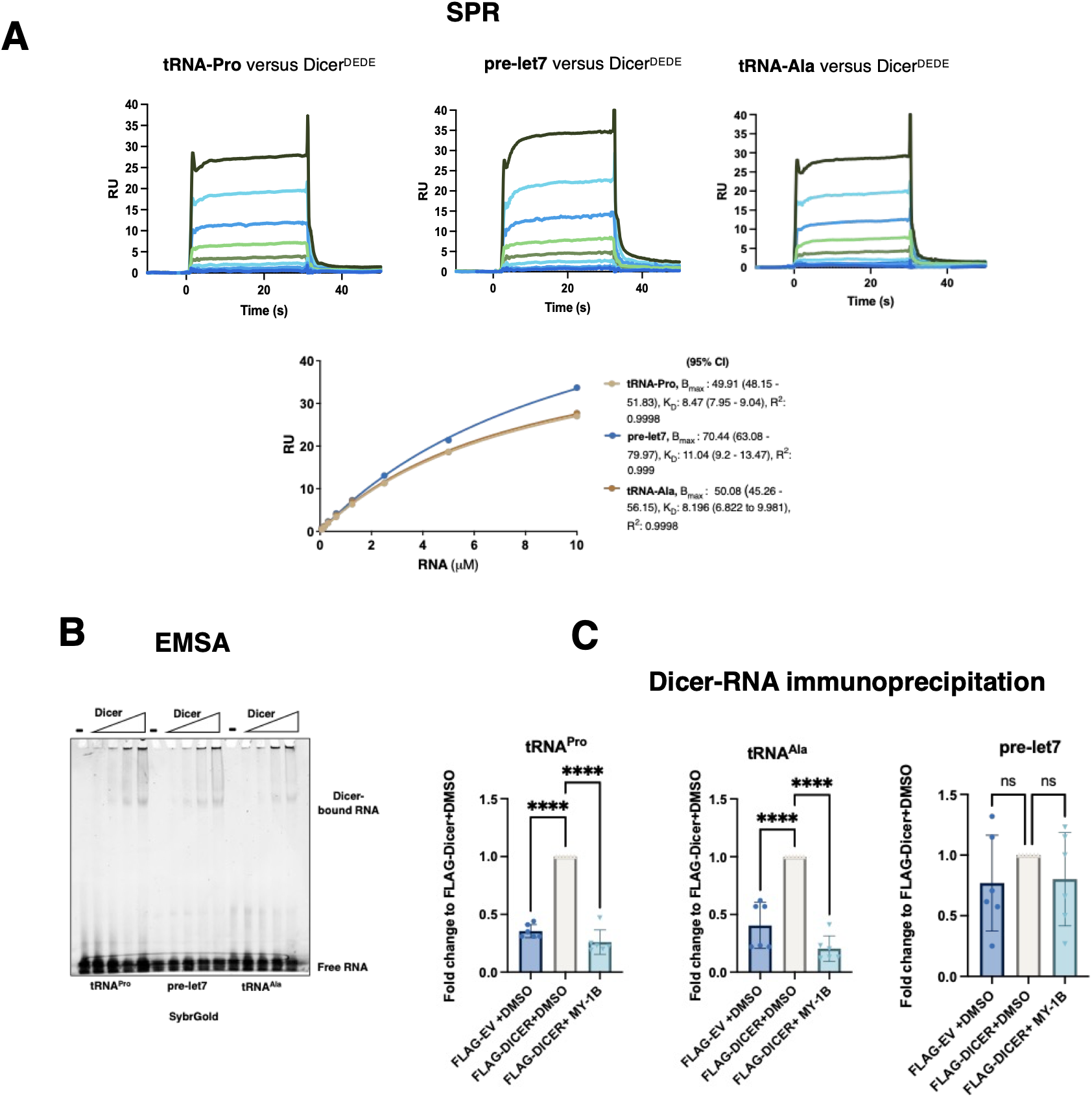
Dicer binds tRNA. **(A)** tRNA-Pro, pre-let7a and tRNA-Ala bind hDicer^DEDE^ with comparable affinity, as measured by surface plasmon resonance (SPR), RU=response units, K_D_ are 8-11 pM. **(B)** Comparative electrophoretic mobility shift assay showing binding of tRNA-Pro, pre-let7a, and tRNA-Ala to hDicer^DED^. **(C)** FLAG-hDicer^WT^ RNA immunoprecipitation from HEK293T cells in WT and treated with NSUN2 inhibitor, showing signals for tRNA-Pro, tRNA-Ala and pre-let7a.

Endogenous tRNA-Pro contains three m^5^C nucleotides at the positions C_47-49_ located near the variable length loop, deposited by RNA methyltransferase NSUN2.^41^ To determine whether NSUN2-dependent methylation affects hDicer binding, we repeated the RIP experiment in cells treated with the NSUN2 methyltransferase inhibitor MY-1B^58^. NSUN2 inhibition significantly reduced the amount of tRNA co-immunoprecipitated with hDicer (Fig. 1C), indicating that m^5^C modification promotes hDicer-tRNA association *in vivo*.

Together, these findings establish tRNA-Pro as an hDicer substrate that binds with affinity comparable to the canonical pre-let7a substrate. They further demonstrate that RNA binding alone does not determine substrate cleavage, consistent with previous reports^59^ and identify NSUN2-dependent methylation as a factor that enhances hDicer-tRNA interaction in the cellular context, suggesting a role for NSUN2 in tsRNA biogenesis.

### hDicer binds NSUN2-methylated tRNA

To further investigate the role of NSUN2, we focused on its interaction with hDicer. We previously showed that hDicer interacts with NSUN2^18^, raising the question of whether this interaction also contributes to tRNA processing. To visualise the interaction between NSUN2 and hDicer in cells, we performed a Proximity Ligation Assay (PLA). This revealed a median of approximately 50 foci per nucleus, with additional foci detected in the cytoplasm, indicating that the interaction between the two proteins extends beyond the nucleus (Fig. 2A). We next investigated the architecture of the NSUN2-hDicer-tRNA complex *in vitro*. Biotinylated hDicer^DEDE^ was immobilized on a SPR sensor chip and binding was measured using purified NSUN2 (Supplementary Fig. S2A), in the presence or absence of the methyl donor S-adenosyl-methionine (SAM) and tRNA-Pro. NSUN2 alone or in combination with either SAM or tRNA-Pro bound hDicer with similar affinities (K_D_ = 0.45-0.62 μM). However, the presence of both SAM or tRNA-Pro resulted in stronger binding, with lower analyte concentrations required to saturate immobilized hDicer (Fig.2B, K_D_=0.45 and 0.52 μM), suggesting that these components stabilize the NSUN2-hDicer complex and likely represent the most biologically relevant assembly (Fig. 2B).

**Figure 2.**
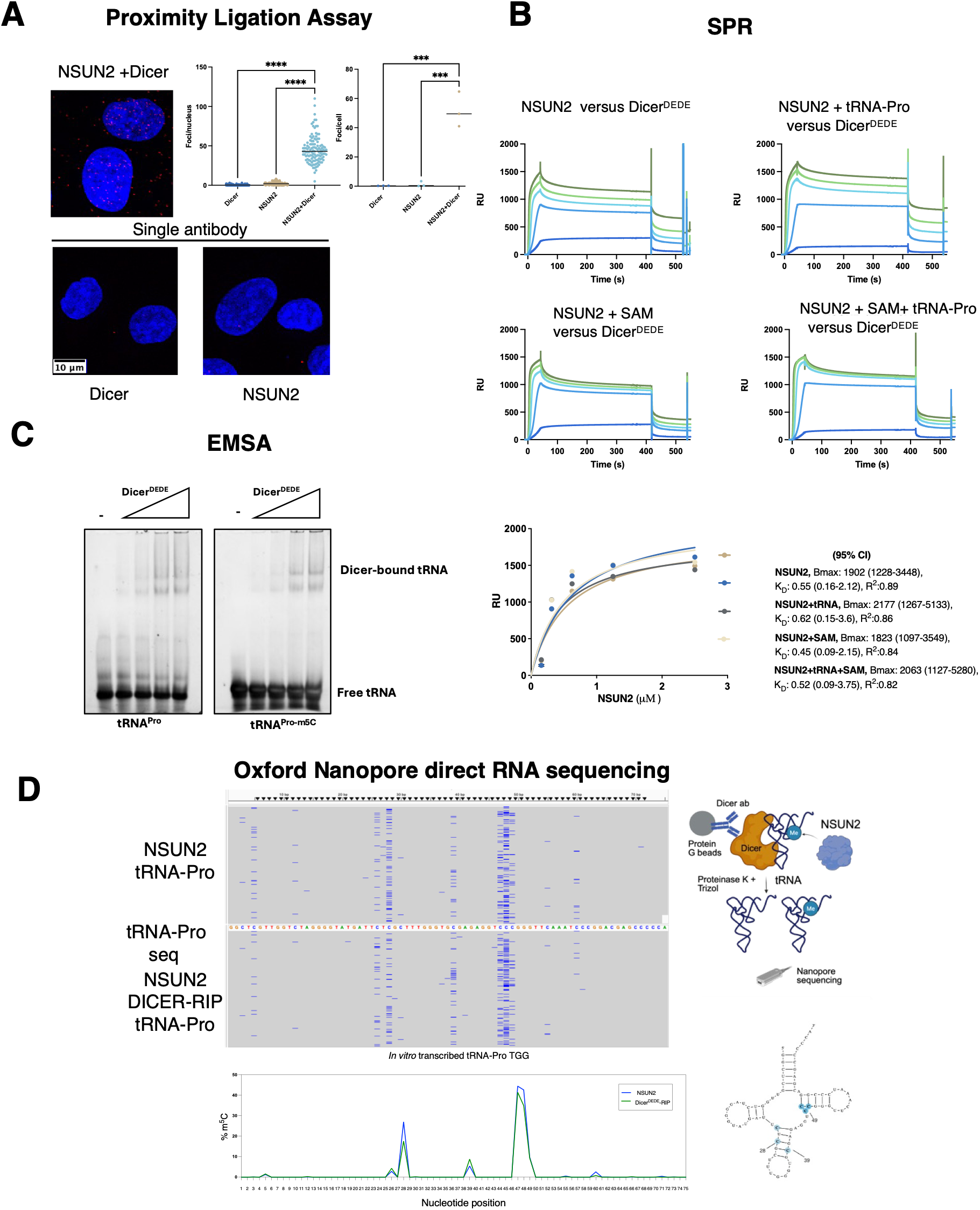
Dicer binds NSUN2 and NSUN2-methylated tRNA. **(A)** NSUN2 and Dicer interact in HeLa cells as shown by Proximity ligation assay, and single antibody controls, showing no unspecific foci. n>100 cells were quantified from n=3 biological repeats. Statistical analysis is Kruskal-Wallis test with Dunn’s multiple comparisons test, ****p-value<0.0001. **(B)** NSUN2 in complex with tRNA and SAM binds hDicerDEDE with higher binding capacity compared to NSUN2, NSUN2 and tRNA or NSUN2 and SAM, as measured by surface plasmon resonance, RU = response units. **(C)** Comparative electrophoretic mobility shift assay showing binding of tRNA-Pro and tRNA-Pro^m5C^ to hDicer^DEDE^ . **(D)** hDicerDEDE binds NSUN2-modified tRNA(m5C), as shown by Oxford Nanopore direct RNA sequencing of samples incubated in vitro with NSUN2 (NSUN2) and hDicerDEDE RNA immunoprecipitation (Dicer RIP). Schematic of in vitro Dicer-RNA immunoprecipitation (RIP) and 2D representation of tRNA-Pro with m5C sites in blue (right).

To test whether hDicer binds NSUN2-methylated tRNA, we synthesized tRNA-Pro containing three m^5^C modifications at positions C47-C49 and performed EMSAs. These experiments confirmed that indeed hDicer binds both methylated and unmethylated tRNA-Pro substrates (Fig. 2C). Finally, to assess whether hDicer preferentially binds methylated or unmethylated tRNA in the presence of NSUN2, we methylated tRNA-Pro *in vitro* using purified NSUN2 and incubated the reaction with hDicer^DEDE^ , followed by hDicer RIP and Oxford Nanopore Direct RNA sequencing (Supplementary Fig. S2B). Approximately 40% of the reads mapping to tRNA-Pro in the NSUN2-treated sample contained m^5^C modification at positions C47-C49 indicating that the *in vitro* methylation reaction did not reach completion under the conditions tested. Notably, tRNA-Pro molecules recovered in the hDicer-RIP also contained these chemical modifications, demonstrating that hDicer associates with methylated tRNA (Fig. 2D). Collectively, these results identify both NSUN2 and methylated tRNA as hDicer interaction partners in both *in vivo* and *in vitro* conditions.

### Cryo-EM of hDicer-tRNA^(m5C)^ complex

To elucidate the structural basis of hDicer recognition of NSUN2-methylated tRNA (tRNA^(m5C)^) by single-particle cryo-EM, we reconstituted the hDicer-tRNA^(m5C)^ complex *in vitro*. *In vitro* transcribed tRNA-Pro was first incubated with NSUN2 to allow methylation and subsequently used to assemble a complex with purified hDicer (Supplementary Fig. S3A). Size-exclusion chromatography revealed co-elution of NSUN2 with the hDicer-tRNA complex, suggesting the formation of a transient trimeric complex, consistent with the SPR results (Supplementary Fig. S3B). Despite this biochemical evidence, cryo-EM analysis did not indetify a particle population corresponding to a stable NSUN2-hDicer-tRNA complex. Instead, extensive particle classification yielded only classes representing the hDicer-tRNA complex and monomeric NSUN2, the latter of which could not be refined to high resolution (Supplementary Fig. S4). Using single-particle cryo-EM, we determined a partial structure of the hDicer-tRNA^(m5C)^ complex at an overall resolution of 3.4 Å (Table1, Fig. 3A, B and Supplementary Figs. S4, S5 and S6). Consensus refinement of the complete particle resolved the conserved L-shape architecture of hDicer, but did not reveal interpretable density for the bound tRNA. Only local refinement focused on the PAZ/Platform domain together with the tRNA improved the density sufficiently to resolve a partial double helical structure (Fig. 3C and Supplementary Fig. S4). Intriguingly, the tRNA density resembles a hairpin structure rather than the canonical cloverleaf structure predicted for tRNA-Pro (Fig. 3D).

**Figure 3.**
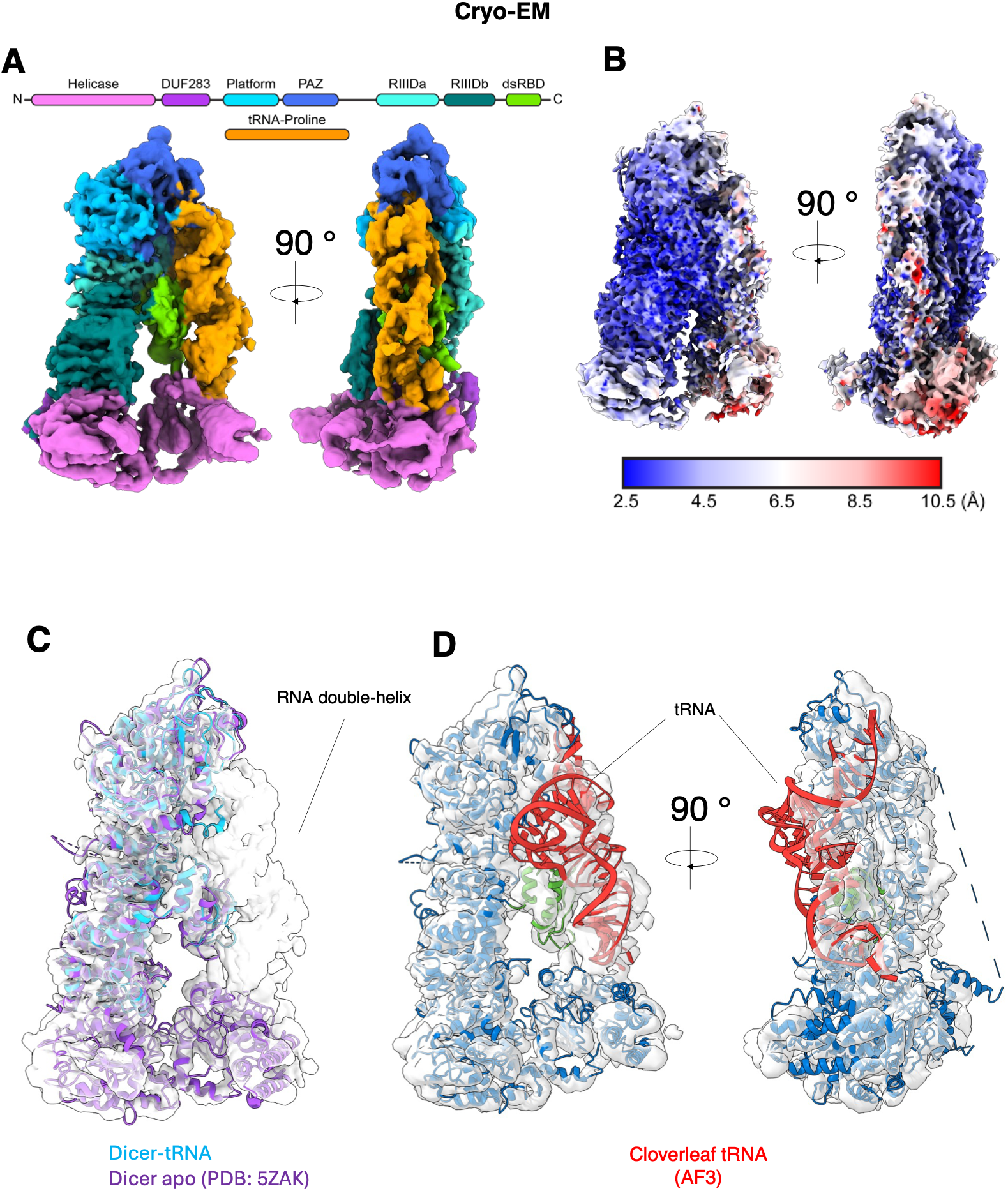
Structure of Dicer-tRNA complex. **(A)** The cryo-EM composite map is coloured by domain structure of Dicer with the tRNA substrate in orange. **(B)** Consensus map coloured by local resolution. **(C)** Structural models of Dicer are fitted in the cryo-EM map: in purple, Dicer in the apo state (PDB ID: 5ZAK), and in blue, built model of hDicerDEDE without the helicase domain. **(D)** AlphaFold 3 model of Dicer with tRNA in a cloverleaf shape is fitted into cryo-EM map with the dsRBD highlighted in green.

Because the observed RNA density adopted a double-helix conformation, typical for miRNA, we considered the possibility that it originated from a contaminating RNA species, such as pre-miRNA co-purified from the Sf9 cells used for protein expression. To exclude this possibility, without a doubt, purified hDicer^DEDE^ was analysed by gel electrophoresis followed by total RNA staining, which detected no contaminating RNA species (Supplementary Fig. S3C). In addition, we sequenced the RNA associated with the purified NSUN2 and hDicer^DEDE^ samples after incubation with tRNA-Pro. Alignment of sequencing reads to the *Spodoptera frugiperda* (Sf9) genome revealed less than 0.1% contaminating RNA, confirming that the density observed in the cryo-EM corresponds to tRNA-Pro (Supplementary Fig. S3D).

To improve the local resolution of the RNA, we tested multiple refinement strategies using masks encopassing the tRNA density together with either the PAZ, dsRBD or platform domains, with different fulcrum positions to account for potential interactions between the tRNA and the respective domains. However, none of these approaches improved the reconstruction, suggesting substantial translational freedom of the tRNA relative to hDicer. This interpretation is consistent with the diffuse tRNA density observed in high-resolution 2D class averages (Supplementary Fig. S4). Furthermore, 3D variability analysis and 3D flex refinement revealed the expected flexibility of the helicase domain but also showcased conjoined movement of the tRNA region, and a modest displacement of the dsRBD (Supplementary Videos 1 and 2). This conformational flexibility is consistent with the previously described transitions between the pre-dicing and dicing states of hDicer, despite the inability of the catalytically inactive hDicer mutant to cleave the RNA substrate^9^.

The final composite reconstruction map, which combines the consensus map, a local refinement excluding the helicase domains and a local refinement of the tRNA/PAZ/Platform region, provided the highest overall map quality (Fig. 3A, C). Although the intrinsically flexible helicase and adjacent DUF283 domains remained poorly resolved, the other regions of hDicer were well defined. The final atomic model (PDB: 9TE0) includes the Platform, PAZ, RNase IIIa and RNAse IIIb domains, together with the dsRBD, whereas the tRNA was not modelled due to the limited local resolution (Fig. 3A-C and Supplementary Fig. S5). Notably, the position of the dsRBD, which undergoes substantial rearrangement during RNA loading into the catalytically active centre^14^, closely resembles that observed in both the apo- and pre-dicing structures of hDicer^13^ (Fig. 3C and Supplementary Fig. S6A and B). In contrast, the positions of pre-miRNA in the previously reported pre-dicing and dicing states do not align with the tRNA tRNA density observed in our reconstruction (Supplementary Fig. S6C and S6D), suggesting that the captured tRNA conformation represents either an average of multiple conformational states or an intermediate along the transition between the pre-dicing and dicing conformations.

### Alternative conformations of tRNA-Pro

The conformation of the tRNA density observed in the cryo-EM map, together with the flexibility at the RNA-protein interface, suggested that tRNA-Pro adopts alternative conformations (Supplementary Video 1). Previous *in silico* studies predicted that zebrafish tRNA-Pro can fold into a hairpin structure ^24^, while *in vivo* Dicer-RIP combined with icSHAPE-MaP identified Dicer-bound tRNAs with non-canonical structures^17^. However, direct structural evidence demonstrating whether these alternative conformations are an intrinsic property of tRNA molecules or are induced upon hDicer binding has been lacking.

To address this question, we performed RNA structure probing using Selective 2′-Hydroxyl Acylation Analyzed by Primer Extension (SHAPE) on free and hDicer-bound tRNA-Pro. In addition, we analysed the secondary structure of tRNA-Alanine AGC 2-1 (tRNA-Ala), a tRNA isotype that is not cleaved by hDicer *in vivo*^23^. All SHAPE experiments were performed using *in vitro* transcribed and re-folded tRNA containing the 3’ CCA trinucleotide. I*n vitro* transcribed tRNAs have been shown to recapitulate the overall architecture of endogenous molecules, including key tertiary contacts^64^.

The tRNAs were probed using either dimethyl sulfate (DMS), which methylates exposed adenine and cytosine residues to generate N¹-methyladenine and N³-methylcytosine ^65^, or the SHAPE reagent 5-Nitroisatoic Anhydride (5NIA), which forms a chemical adduct (2’-O) at the 2′ hydroxyl group of an accessible ribose via ester linkage^66^. Reactivity values were normalised to DMSO-treated controls and used as experimental restraints for secondary structure prediction using the ViennaRNA package^67^. Reproducible reactivity profiles were obtained across biological replicates and both chemical probing methods for all RNA species analysed (Supplementary Fig. S7A-F, S8A-D). RNA structure probing of tRNA-Ala yielded the expected canonical cloverleaf secondary structure, comprising four short dsRNA stems and three ssRNA loops, corresponding to the anticodon, D (dihydrouridine), TΨC (Thymine-Pseudouridine-Cytosine) arm/loop and the acceptor stem (Supplementary Fig. S7C and F). In contrast, SHAPE guided prediction of tRNA-Pro revealed a markedly different conformation. Both free and hDicer-bound tRNA-Pro adopted a hairpin-like structure containing a prominent bulge and an extended stem rather than the canonical cloverleaf fold (Fig. 4A-C and Supplementary Fig. S8A-D). Comparison of alternative tRNA-Pro models identified a flexible region spanning nucleotides 8-21 that can either remain single stranded or partially base-pair to form a short five-base-pair stem and a smaller bulge (Fig. 4B). Upon hDicer binding this same region exhibited reduced chemical reactivity, suggesting local stabilization of the flexible segment, analogous to the conformational stabilization previously observed during pre-miRNA recognition by hDicer^13^ (Fig. 4C and Supplementary Fig. S8, S9A).

**Figure 4.**
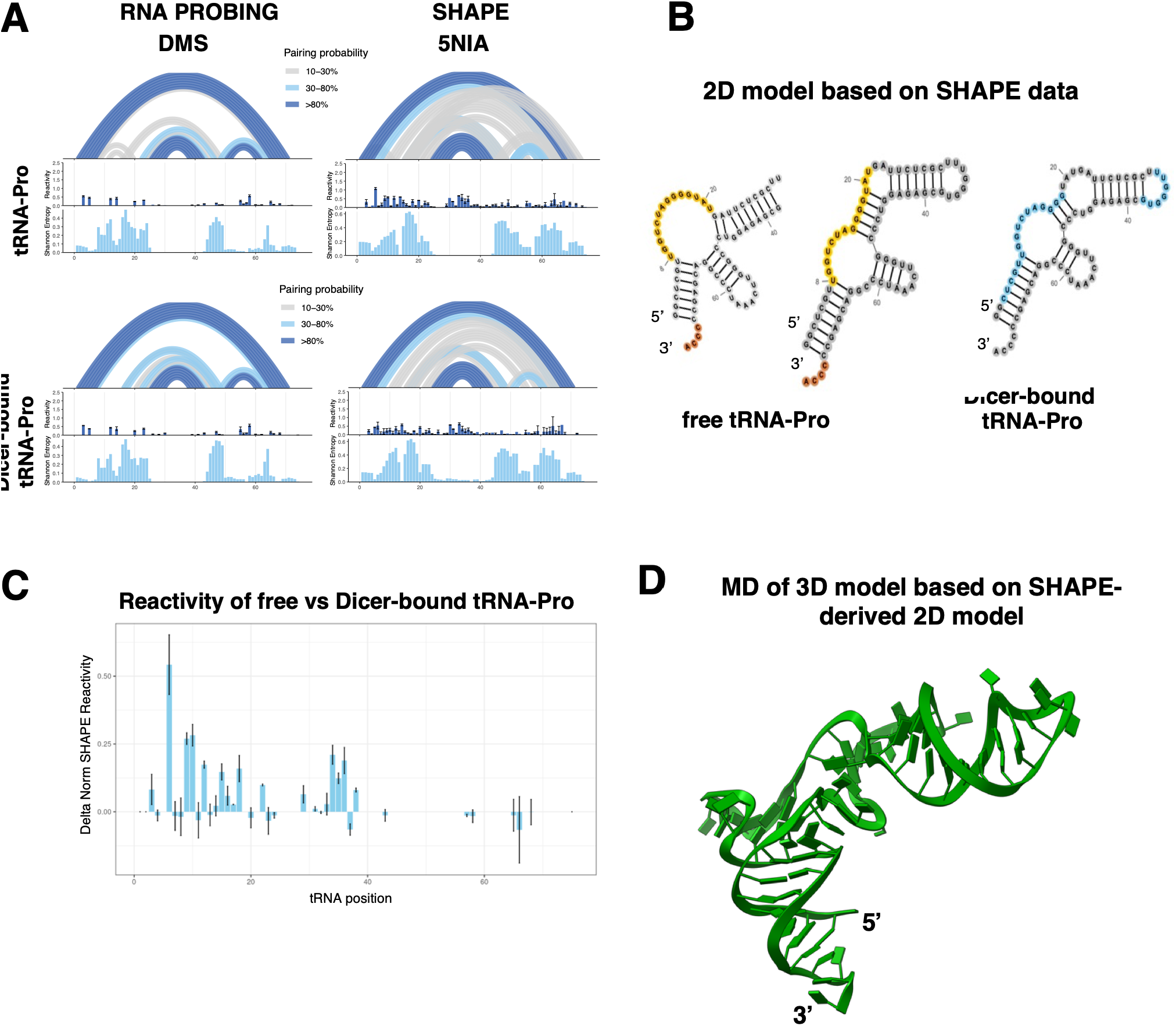
tRNA-Pro folds into alternative conformations. **(A)** SHAPE reactivity and Shannon entropy profiles of free and Dicer-bound tRNA-Pro of combined replicates (n=3). **(B)** Alternative conformations of free tRNA-Pro (left and middle) and Dicer-bound tRNA-Pro (right), highlighting flexible (orange) and regions affected by changes in reactivity upon Dicer binding (blue). **(C)** Changes in reactivity of free versus Dicer-bound tRNA-Pro, showing overall higher reactivity of free versus Dicer-bound tRNA-Pro (also see **Supplementary** Fig. 7 and 8**). (D)** Tertiary structure of free tRNA-Pro. 3D tRNA model was obtained with trRosettaRNA guided by SHAPE at equilibrium, as shown with MD simulations, 1 ps (n=2).

To visualize the three-dimensional organization of the hDicer-bound RNA, we generated a structural model of tRNA-Pro using trRosettaRNA guided by the SHAPE data derived from the hDicer-bound state. The resulting model fitted well into the Dicer-tRNA cryo-EM density, illustrating how the flexible region of tRNA-Pro could be stabilised through contacts with the dsRBD of hDicer (Supplementary Fig. S9A, B). Because protein binding can reduce nucleotide accessibility to SHAPE reagents and thereby bias structural inference, we independently modelled free tRNA-Pro using the SHAPE-derived secondary structure obtained in the absence of hDicer (Fig. 4D). This model was subsequently subjected to molecular dynamic (MD) simulations. Following initial equilibration, the tRNA conformation relaxed into a more solvent exposed conformation while preserving its overall secondary structure throughout the simulation trajectory (Supplementary Fig. S9C). These simulations indicate that the hairpin like fold is structurally stable yet conformationally flexible (Fig. 4D).

Overall, the cryo-EM, SHAPE and MD analyses support the existence of a stable non-canonical tRNA-Pro conformation, that is intrinsically adopted in solution and recognised by hDicer.

### Architecture of the hDicer-tRNA complex

To characterise the architecture and dynamics of the hDicer-tRNA complex, we fitted the 3D model of tRNA-Pro into our cryo-EM map and performed MD simulations. Throughout the simulation, the complex remained stable without major unfolding or disruption of either the protein or the RNA. Notably, the tRNA underwent a conformational rearrangement, forming closer contacts with the PAZ domain and adapting to the surface of hDicer, in agreement with the cryo-EM density (Fig. 5A and Supplementary Fig. S9D and S9E). The central part of the tRNA forms extensive interactions with the dsRBD and the RNAase domains, rich in arginine and lysines, while the apical loop (hairpin) region of the tRNA exhibits high flexibility as indicated by the RMSF plots (Supplementary Fig. S9E). This flexibility is consistent with observed changes in RMSD, which reflect the conformational rearrangement of the tRNA as it shifts closer to the protein (Supplementary Fig. S9E). Consistent with the SHAPE experiments, the MD simulations also predicted reduced accessibility of nucleotides that became buried upon hDicer binding. Overall, the MD-refined model remained stable throughout the trajectory, improved the fit to the cryo-EM density, and provides a structural model for the experimentally observed hDicer–tRNA complex (Supplementary Fig. S9D and E).

**Figure 5.**
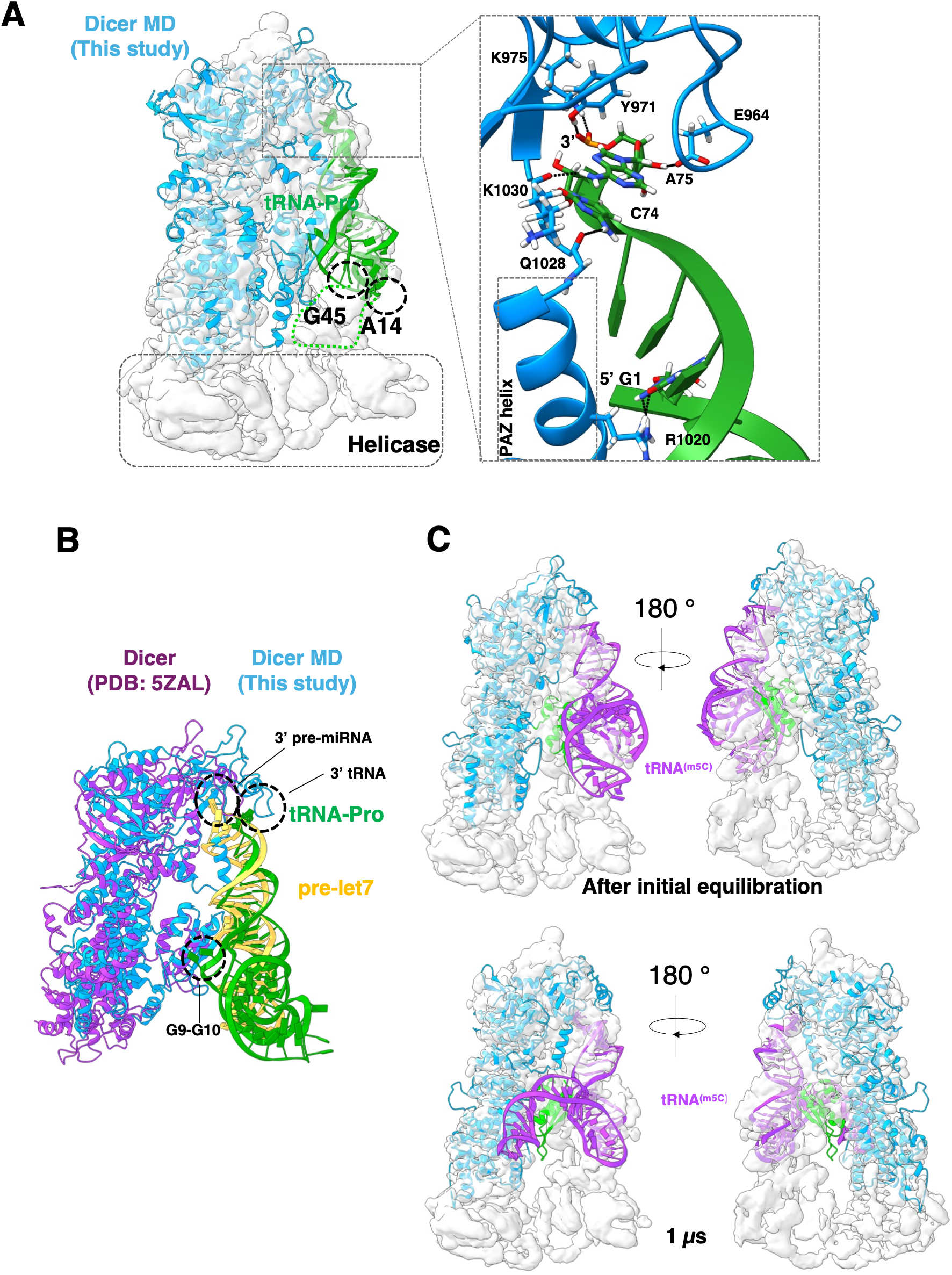
Molecular dynamics reveals a model for tRNA-Pro and tRNA-Pro^m5C^ bound by Dicer. **(A)** MD model of Dicer-tRNA-Pro complex fitted in the cryo-EM map, and magnified view of Dicer PAZ domain anchoring the 3’ end of the tRNA. **(B)** Structural comparison of our model of Dicer-tRNA (replica 1) with Dicer in complex with pre-let7 (PBD:5ZAL). **(C)** tRNA<^m5C^> promotes Dicer cleavage by inducing conformational change from ’close’ to ’open’ state similar to pre-dicing to dicing transition, as shown by molecular dynamics simulations, after initial equilibration (left) and 1 ps (right) *(n=2).*

In the refined complex, the acceptor stem together with portions of the D and TΨC arms (G1– A14 and G45–A75) forms an extended double-stranded RNA helix (Fig. 5A). The terminal 3′ nucleotide (A75) establishes potential hydrogen-bonding interactions with residues Y971, E964, K975 and K1030 of hDicer, whereas nucleotide C74 may interact with Q1028 (Fig. 5A). When compared to hDicer in complex with canonical pre-let7 substrate (PDB:5ZAL)^13^, the tRNA-Pro arms align to pre-let7, but while pre-let7 is a straight hairpin, a region of tRNA-Pro that assembles into the ssRNA central loop, nucleotides G9 and G10, deviates from the dsRNA helix and may be contacted by arginine residues (such as R1907 and R1910) of the dsRBD potentially through hydrogen bonding, corroborating the SHAPE data (Fig. 5B). When comparing the ends of the two RNA substrates, unlike the pre-miRNA, the 5′ end of the tRNA G1 establishes a potential contact with R1020, a residue located in the human PAZ helix, while the 3′ end of both RNA molecules are anchored by the PAZ domain involving contacts with Y971 shared by both substrate, the pre-miRNA inserts deeper into the PAZ 3′ pocket ^13,14^ compared to tRNA that instead may be also contacted by residues in the PAZ helix (Fig. 5A, B). Interactions between the apical anticodon loop are not represented in this model, as we excluded the helicase domain from the molecular dynamic simulations, due to the flexibility of this region. Together, these analyses provide the first structural model of hDicer bound to a non-canonical tRNA substrate.

Despite detecting binding of tRNA^(5mC)^ by hDicer (Fig. 2C, D), we performed SHAPE with unmethylated tRNA to investigate structural differences of tRNA isotypes solely based on nucleotide sequence. To identify the structural basis of tRNA^(5mC)^ cleavage by hDicer, here we introduced m^5^C in place of the three cytosine C47, 48 and 49 in tRNA-Pro *in silico* and simulated the molecular dynamics of the Dicer-tRNA^(m5C)^ complex. Indeed, this model showed that m^5^C_47-49_ promotes an alternative conformation of the Dicer-bound tRNA, which can partially displace the dsRBD (Fig. 5C, Supplementary Fig. S10 A and B and Supplementary Video 3). This allows the tRNA to get closer to hDicer, compared to the unmodified tRNA, which could facilitate the pre-dicing to dicing transition^14^ (Supplementary Fig. S10). These structural rearrangements are likely to promote cleavage of tRNA^(m5C)^ and corroborate the flexibility around the dsRBD observed in our structural analyses. This together with the direct RNA sequencing of hDicer^DEDE^-RIP, suggest that we may have captured conformations of hDicer transitioning between states, potentially in complex with unmodified tRNA and tRNA^(m5C)^ (Supplementary Video 2).

Collectively, the MD simulations demonstrate that hDicer forms a stable yet highly dynamic complex with tRNA and provide a molecular model that integrates the cryo-EM and SHAPE data. The flexibility of both the protein and the RNA enables the tRNA to adapt to multiple hDicer domains during substrate recognition, whereas m5C modification promotes conformational rearrangements that favour progression towards the catalytically competent dicing state.

### NSUN2 facilitates hDicer-cleavage of tRNA

Several hDicer interacting proteins, including ADAR-1 and TRBP, enhance the cleavage efficiency of canonical hDicer substrates ^68,69^. We therefore investigated whether NSUN2 similarly promotes hDicer-mediated processing of tRNA. To validate the molecular dynamics (MD) predictions that m5C modification facilitates hDicer cleavage, we performed in vitro methylation of tRNA-Pro using purified NSUN2, followed by cleavage assays with purified wild-type hDicer (Supplementary Fig. S11A). Catalytically inactive NSUN2^K190M and hDicer^DEDE^ mutants were included as controls. Indeed, pre-incubation of the tRNA-Pro with NSUN2 and SAM, allowing for tRNA^(m5C)^, significantly increased hDicer cleavage efficiency (Fig. 6A). In contrast, this was reduced in presence of NSUN2 K190M or hDicer^DEDE^ mutants demonstrating that both NSUN2 methyltransferase activity and hDicer catalytic activity are required (Fig. 6A).

**Figure 6.**
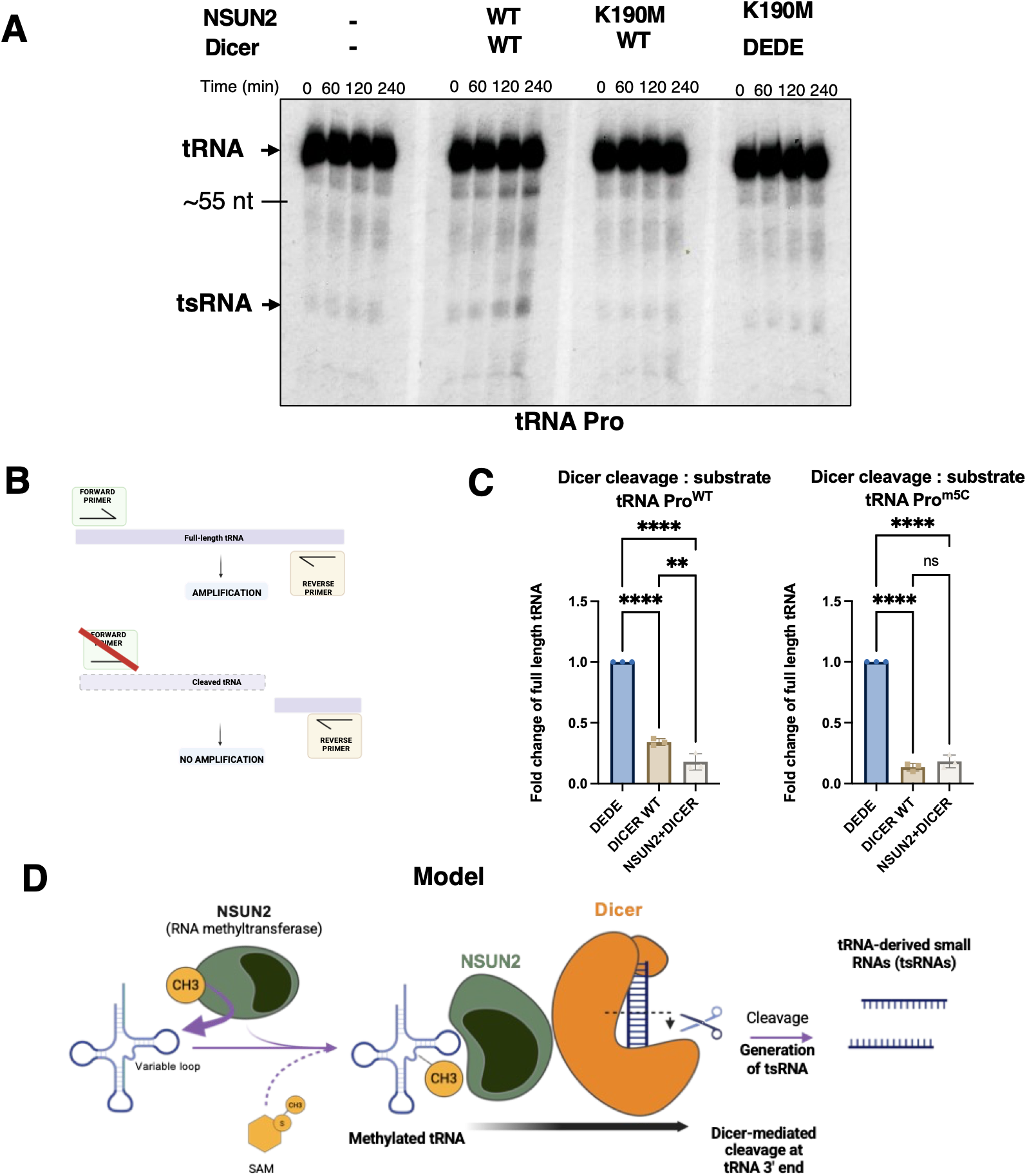
NSUN2 facilitates Dicer cleavage of tRna-Pro. (A) Representative Northem blot of tRNA-Pro cleaved by Dicer after methylation by NSUN2 WT or incubation with methylation-deficient mutant NSUN2^K190M^ and inactive mutant Dicer^DEDE^. (B) Schematic of qRT-PCR for quantification of cleaved tRNA (c) Quantification of full-length tRNA-Proc^m5C^ substrate after cleavage by Dicer, (*n=3*). (D) Schematics of model of NSUN2-Dicer interplay in tsRNA biogenesis. Image generated with BioRender.com

Because Northern blotting provides only semi-quantitative measurements, we next quantified the remaining full-length tRNA substrate by RT-qPCR following the cleavage reaction (Fig. 6B). As expected, hDicer significantly reduced the abundance of full-length tRNA-Pro, and this effect was further enhanced by NSUN2-mediated methylation.

Our direct RNA sequencing experiments showed that the in vitro NSUN2 methylation reaction modified approximately 40% of tRNA molecules (Fig. 2D). To determine whether complete methylation further enhances cleavage, we compared hDicer activity on chemically synthesized tRNA-Pro containing m5C modifications at positions C47–C49 with that on the corresponding unmethylated substrate. The methylated tRNA was cleaved more efficiently (approximately 90%) than the unmethylated substrate (approximately 70%), reaching a level comparable to that observed for unmethylated tRNA pre-incubated with NSUN2 (approximately 80%) (Fig. 6C). Furthermore, pre-incubation of the fully methylated synthetic tRNA-Pro(m5C) with NSUN2 did not further increase cleavage efficiency, indicating that m5C modification itself is sufficient to account for the stimulatory effect (Fig. 6C).

Finally, to investigate whether m5C influences tRNA conformation independently of hDicer, we folded unmethylated and methylated tRNA-Pro in vitro and analysed them by native polyacrylamide gel electrophoresis. Whereas unmethylated tRNA-Pro resolved into multiple conformational species, tRNA-Pro(m5C) migrated predominantly as a single species, indicating that m5C stabilizes a specific tRNA conformation (Supplementary Fig. S11B).

Collectively, these results demonstrate that NSUN2-mediated m5C modification stabilizes tRNA structure and directly promotes its efficient cleavage by hDicer, providing experimental validation of the structural mechanism predicted by the MD simulations.

## DISCUSSION

tsRNA molecules have increasingly been implicated in diverse cellular processes, making them important molecular regulators ^27–37,70^. Beyond their biological roles, tsRNAs are also being explored as translational and therapeutic assets, including as biomarkers of multiple cancers and other diseases ^27–36^ and as oligonucleotide tool for nuclear gene silencing ^23^. As efforts to define tsRNA biogenesis and regulation have progressed, hDicer has emerged as one enzyme contributing to the generation of specific tsRNA subsets,^20–25^, including fragments derived from the 3′ and 5′ ends of tRNA molecules^23^.

Although hDicer recognition^7,10–12^, binding^13,17^ and cleavage of canonical substrates, particularly pre-miRNA,^7,14^ have been extensively characterized by structural ^9,13,14^ and biochemical approaches^10–12^, the molecular basis governing hDicer processing of non-canonical substrates remains incompletely understood. Cellular studies have detected tRNAs bound to hDicer^17,59^ but whether these molecules are converted into functional products by hDicer was not resolved. In earlier work, we provided direct evidence that hDicer can process tRNAs^23^, which are structurally distinct from canonical Dicer substrates, such as hairpin RNAs. Here, we visualised the complex of human Dicer with a non-canonical tRNA substrate, using SHAPE probing, cryo-EM and MD simulations. The resulting model indicates that, upon binding hDicer, the tRNA adopts a dsRNA helix structure. Notably, when compared with the pre-miRNA-hDicer complex^13^, the overall architecture is highly similar (Fig. 5B). In our model, the tRNA is anchored its 3′ end by the PAZ domain and engages the dsRBD, which likely contributes to substrate recognition and to positioning the tRNA within the catalytic valley as the complex transitions from a pre-dicing to dicing state.

A key unresolved question has been why hDicer, despite binding, processes only a subset of tRNA isotypes ^23^. Although prior work suggested that some tRNAs may transiently adopt Dicer-compatible, hairpin-like conformations^17,24^, direct evidence for such alternative folding states and their relationship to substrate selectivity has been limited. Using SHAPE RNA probing, we confirm that tRNA can indeed fold into hairpin-like conformations. Importantly, by studying unmodified, *in vitro* transcribed tRNAs, we demonstrate that primary sequence alone can support folding into these non-canonical secondary structures, independent of cellular modification patterns. Consistent with this framework, tRNA-Pro, previously identified as an hDicer substrate, does not predominantly adopt the canonical cloverleaf in our probing experiments, instead it folds into alternative conformations that can be recognized by hDicer and may be further stabilized by the enzyme prior to processing (Fig. 4C). In contrast, tRNA-Ala, which is not processed by hDicer *in vivo* ^23^, folds robustly into the canonical cloverleaf architecture. Moreover, SHAPE reactivity of hDicer-bound tRNA-Pro is globally reduced relative to free tRNA (Fig. 4C). While reduced reactivity is consistent with protein shielding and limited reagent accessibility, it may also reflect hDicer-imposed structural remodelling, potentially mediated by the dsRBD and additional domains such as the helicase, analogous to rearrangements described for pre-miRNA substrates^13^. In canonical processing, such remodelling occurs while the helicase domain occludes access to the catalytic core and constrains progression to the dicing state^9,13^. In our cryo-EM reconstruction, the helicase region is flexible and poorly resolved, and it was excluded from the MD simulations. This limitation likely explains the mobility observed for the tRNA apical loop in our model and the replicate-to-replicate variability across MD trajectories: interactions that would normally restrain this region via contacts with the helicase are not represented. We therefore propose that the helicase domain plays a substantive role in stabilizing and/or guiding tRNA conformations that are competent for cleavage. More broadly, we hypothesize that intrinsic structural plasticity is a prerequisite for hDicer processing: tRNAs that sample alternative conformations in solution can be captured and remodelled by hDicer into a cleavage-competent state. Conversely, tRNAs with particularly stable, well-defined secondary structures-exemplified by tRNA-Ala-may resist the conformational rearrangements required for productive engagement and are therefore excluded from the effective substrate repertoire. In cells, tRNA molecules, are extensively chemically modified to promote maturation, folding and functional accuracy. Several studies have linked RNA modifications to non-coding RNA processing^39–41,71,72^, raising the possibility that RNA modifying enzymes modulate hDicer activity and, by extension, tsRNA biogenesis. In this study, we have identified a functional interplay between NSUN2-dependent cytosine-5 methylation of tRNA and hDicer-mediated processing of tRNA, and we propose NSUN2 as a novel regulator of Dicer in the context of tsRNA biogenesis. However, key aspects of this regulatory pathway, remains to be elucidated, including the cellular site(s) at which NSUN2–Dicer coordination occurs during tsRNA production. Proximity ligation assays detect NSUN2–Dicer association in both nuclear and cytoplasmic compartments (Fig. 2A). Because NSUN2–Dicer interactions have also been implicated in processes not directly related to tsRNA biogenesis ^18^, these observations remain compatible with additional, as-yet-uncharacterized roles, including potential effects on other non-coding RNA pathways.

Size exclusion chromatography indicated co-elution of NSUN2 with the hDicer–tRNA complex, yet we did not observe a stable trimeric NSUN2/hDicer/tRNA assembly in the cryo-EM sample. This suggests that the ternary complex is either transient or weak under the conditions tested. One plausible interpretation is that NSUN2 dissociates rapidly after methyl transfer, consistent with limited and short-lived contacts with hDicer. Supporting this view, MD simulations of methylated tRNA (tRNA^m5C^) bound to hDicer, together with cleavage assays (Fig. 6), indicate that the enhanced processing effect can be attributed to the m^5^C mark on the RNA substrate itself rather than to the persistent presence of NSUN2. We therefore propose a model in which NSUN2–hDicer association can function as a substrate-handover mechanism (Fig. 6D). That said, direct protein–protein interactions should not be excluded: surface plasmon resonance (SPR) supports direct binding in vitro (Fig. 2B), and NSUN2 may contact hDicer, potentially via the helicase domain, consistent with how other hDicer cofactors engage the enzyme^4,13,68^, to promote conformational states favourable for catalysis.

While links between RNA chemical modifications and RNA processing have been reported, the structural basis for this crosstalk has remained largely unexplored. Although inclusion of NSUN2 in our in vitro reconstitution may have increased sample heterogeneity, it enabled us to interrogate how methylation affects hDicer processing of tRNA substrates. Our structural model, together with biochemical measurements (Fig. 3 and 6), suggests that NSUN2-mediated m^5^C modification enhances hDicer processing by promoting conformational changes consistent with the pre-dicing to dicing transition previously described for canonical substrates^14^. The magnitude of cleavage enhancement we observe for NSUN2 is more modest than that reported for established hDicer cofactors such as TRBP and ADAR-1, which can increase activity by several-fold^68,69^. This difference may reflect substrate dependence. Notably, TRBP has been reported to preferentially enhance cleavage of suboptimal substrates and can even inhibit processing in certain contexts^69^. In this light, tRNA-Pro may already represent a relatively optimal hDicer substrate and therefore show only limited additional benefit from methylation. By contrast, other, currently unidentified suboptimal tRNA substrates may exhibit a stronger dependence on NSUN2-mediated modification for efficient processing.

In summary, this work provides, to our knowledge, the first structural model of human Dicer bound to a non-canonical tRNA substrate. Collectively, our findings broaden the repertoire of RNAs processed by hDicer and demonstrate how RNA chemical modifications can tune the biogenesis of small RNAs such as tsRNAs, molecules with substantial diagnostic and therapeutic potential.

## ACKNOWLEDGEMENTS

This work was supported by the Medical Research Council [BVR02590], EPA Trust Fund [BVR01670], and Lee Placito Fund awarded to M.G; the Center of Excellence for HPC H2020 European Commission, BioExcel-3: Centre of Excellence for Computational Biomolecular Research [European Union: 101093290], the MDDB project [101094561], Instituto de Salud Carlos III─Instituto Nacional de Bioinformatica, Fondo Europeo de Desarrollo Regional [ISCIII PT 17/0009/0007], and European Regional Development Fund, ERFD Operative Programme for Catalunya, the Catalan Government AGAUR [SGR2021 00863]. The IRB Barcelona is the recipient of a Severo Ochoa Award of Excellence from MINECO. Modesto Orozco is an ICREA Academia Fellow. Federica Battistini is a Serra Hunter Fellow.

We would like to thank David LV Bauer for his support and help with SHAPE experiments, Matthew K. Higgins for his help with cryo-EM, Ervin Fodor for helpful feedback on manuscript and Mikhail Kutuzov for assistance with SPR.

## AUTHOR CONTRIBUTIONS

A.D.F. designed and performed most of the experiments and prepared initial draft of the manuscript. S.H. performed Cryo-EM analyses and edited the manuscript. F.B. performed MD simulations. N.B.S. assisted with SHAPE experiments and J.B. performed bioinformatic analyses of SHAPE data. K.A. performed the bioinformatic analysis of tRNA Nanopore sequencing. Ab.Ab. created a model of tRNA. A.A. contributed to the original idea and provided help with several technical aspects. M.O. provided expertise and advice on MD simulations. M.G. designed and supervised the project and edited the manuscript. All the authors reviewed and approved the final version of the manuscript.

## SUPPLEMENTARY DATA

Supplementary Data and videos are available at NAR online.

## DATA AVAILABILITY

Cryo-EM maps of the hDicer-tRNA complex are available at the Electron Microscopy Data Base (EMDB) under the accession codes EMD-55806 (consensus map), EMD-55807 (focused map without helicase domain), EMD-55808 (focused map tRNA + PAZ/Platform domains) and EMD-55809 (composite map). The protein model for hDicer (without tRNA) is available at the Protein Data Base (PDB) under the accession code 9TE0.

MD simulations are available at MDposit: https://mmb-dev.mddbr.eu/#/id/A026J/overview https://mmb-dev.mddbr.eu/#/id/A025U/overview https://irb-dev.mddbr.eu/#/id/A026O/overview https://irb-dev.mddbr.eu/#/id/A025V/overview https://irb-dev.mddbr.eu/#/id/A026I/overview

SHAPE and RNA-sequencing data are available in SRA with accession number PRJNA1418148. Codes and the tRNA FASTA files used for the alignment can be found on GitHub: https://github.com/jamesboot/shape/tree/main

