## Supplementary files for "Structural and functional basis of the non-canonical human Dicer-tRNA complex"

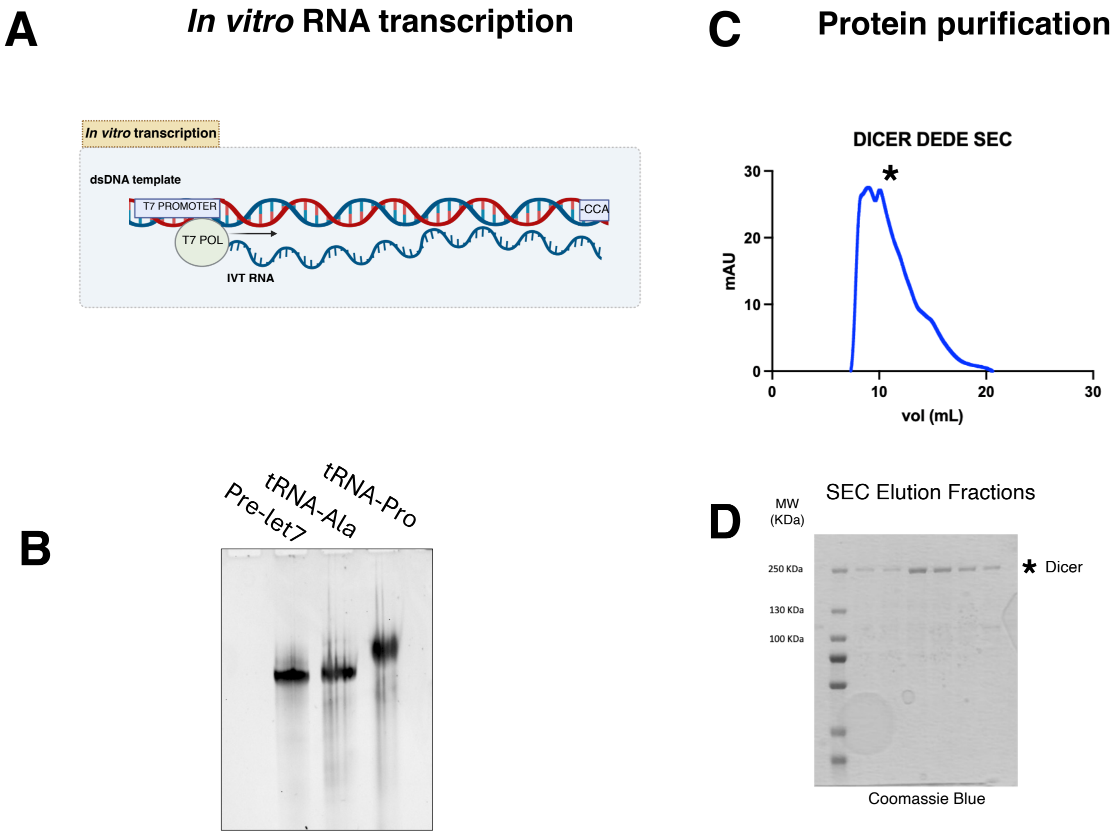


**Supplementary Figure S1.** (**A**) Schematic of *in vitro* RNA transcription. (**B**) Denaturing RNA PAGE showing RNA substrates obtained by *in vitro* transcription. (**C**) Size exclusion chromatography profile and (**D**) reducing SDS-PAGE gel of purified hDicer$^{DEDE}$.

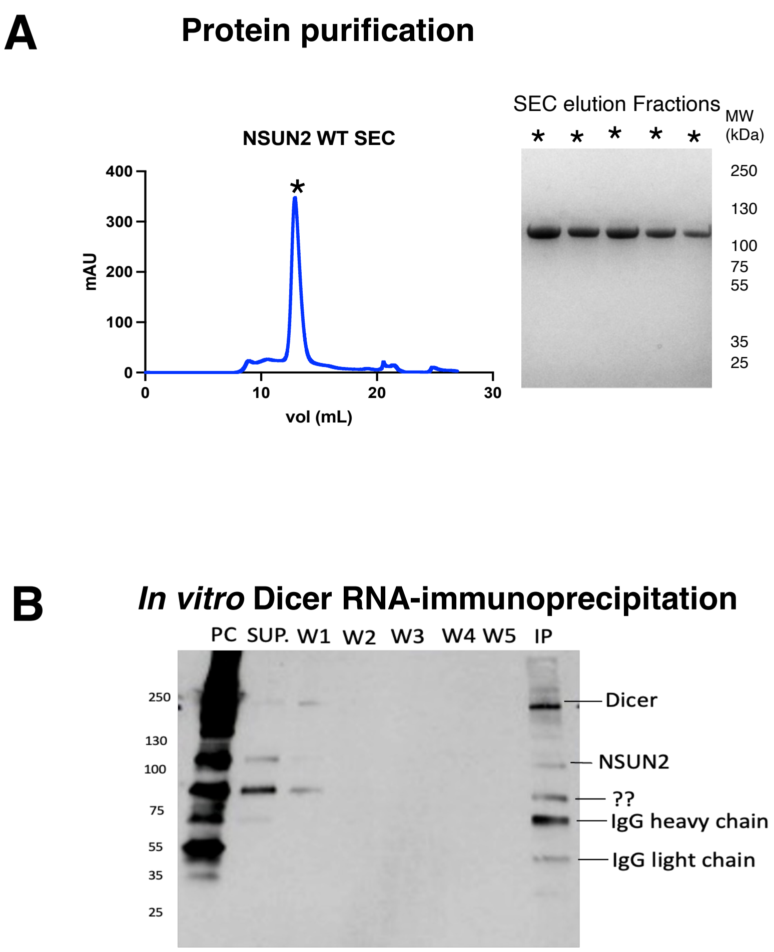


**Supplementary Figure S2.** (**A**) Size exclusion chromatography profile and reducing SDS-PAGE gel of purified NSUN2. (**B**) Western blot of pre-clearing (PC), supernatant (SP), washes (W1-5), and elution (IP) fractions of in vitro Dicer-RIP.

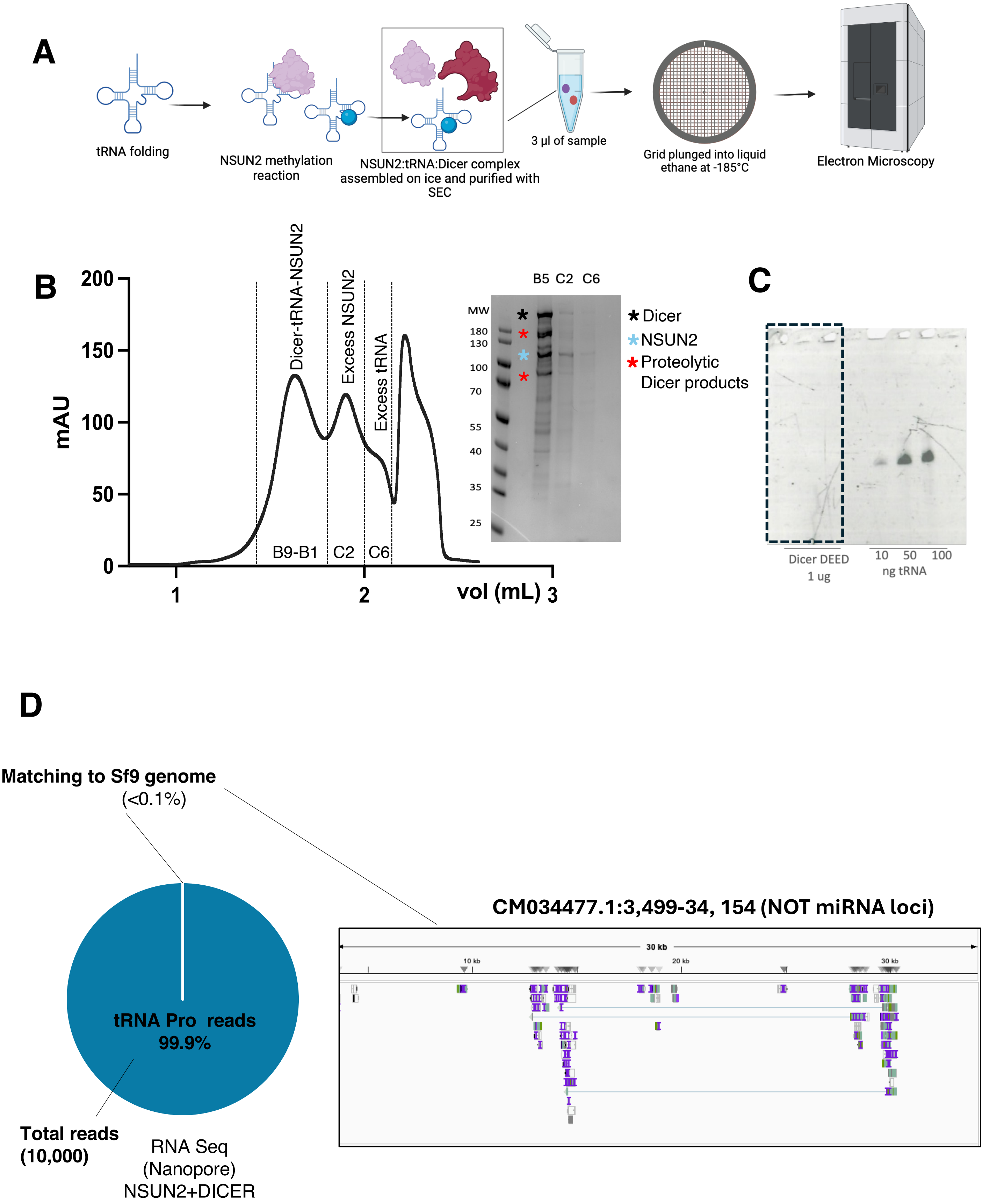


**Supplementary Figure S3.** **(A)** Schematic of *in vitro* reconstruction of the complex for cryo-EM. **(B)** Size exclusion chromatography profile and reducing SDS-PAGE gel of Dicer-tRNA$^{(m5C)}$ complex. **(C)** Denaturing urea-PAGE gel of purified hDicer$^{DEDE}$ and control RNA (10-100 ng). **(D)** Sequencing alignment of DicerDEDE and NSUN2 purified proteins + tRNA-Pro to the genome of insect cells Sf9 used for expression of the recombinant proteins.

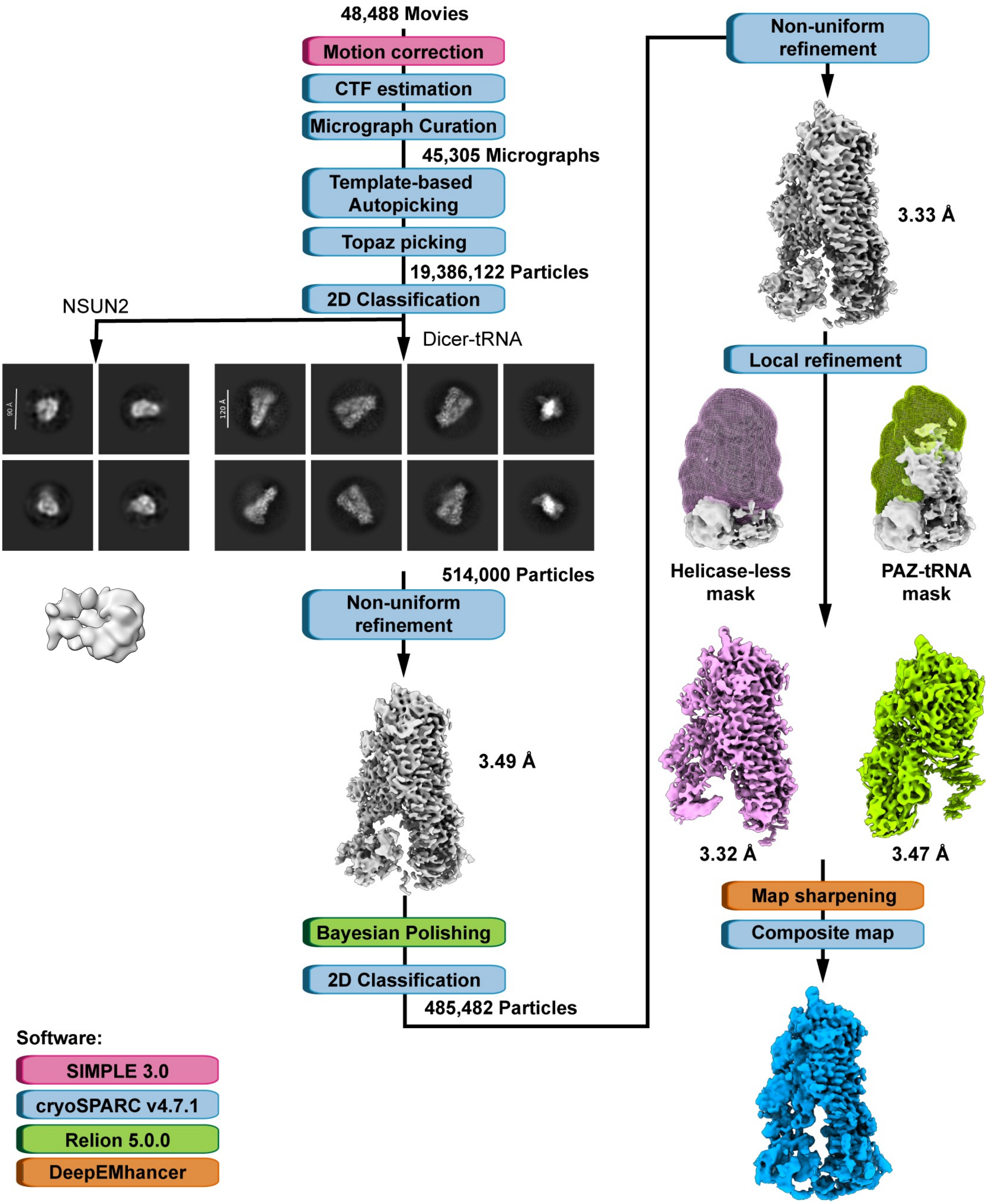


**Supplementary Figure S4.** Single particle cryo-EM data processing workflow. Micrograph movies from two grids were pre-processed in SIMPLE and cryoSPARC, followed by template-based and TOPAZ particle picking in cryoSPARC. Particles were curated by removal of duplicates and multiple rounds of 2D classification, subjected to Bayesian polishing in Relion and non-uniform refinement in cryoSPARC to generate a consensus map of the whole particle. To improve individual regions of the map, it was locally refined using soft masks around the regions of interest. Finally, maps were sharpened using DeepEMhancer and a composite map was generated in ChimeraX using the consensus and the locally refined maps.

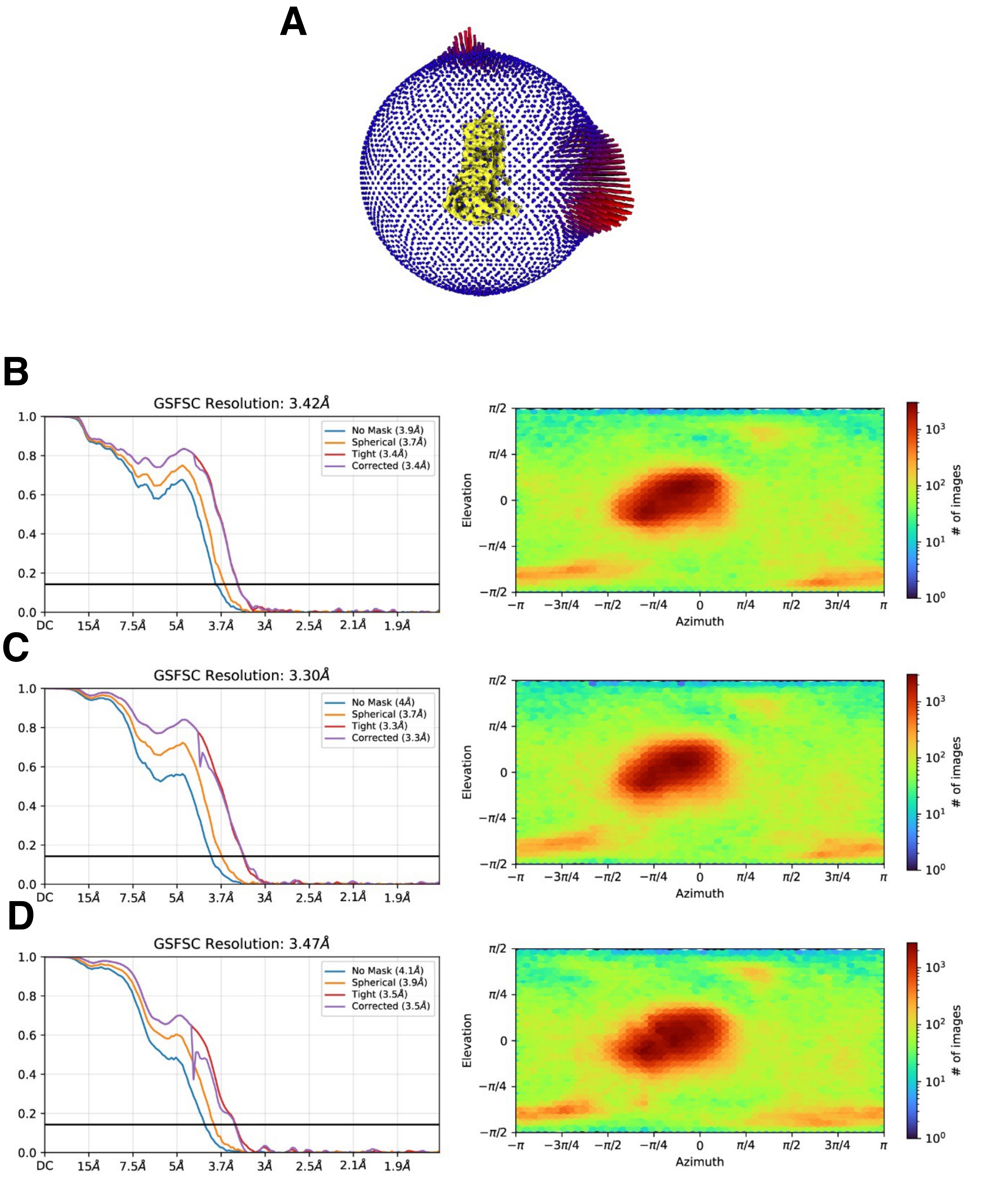


**Supplementary Figure S5.** Cryo-EM map validation and statistics. (**A**) 3D representation of particle Euler angles. (**B**) Gold-standard fourier shell correlation (FSC) curves and particle view distribution of the consensus reconstruction, (**C**) local refinement using a helicase-less mask or (**D**) a mask focused on the PAZ/Platform domains and the tRNA region.

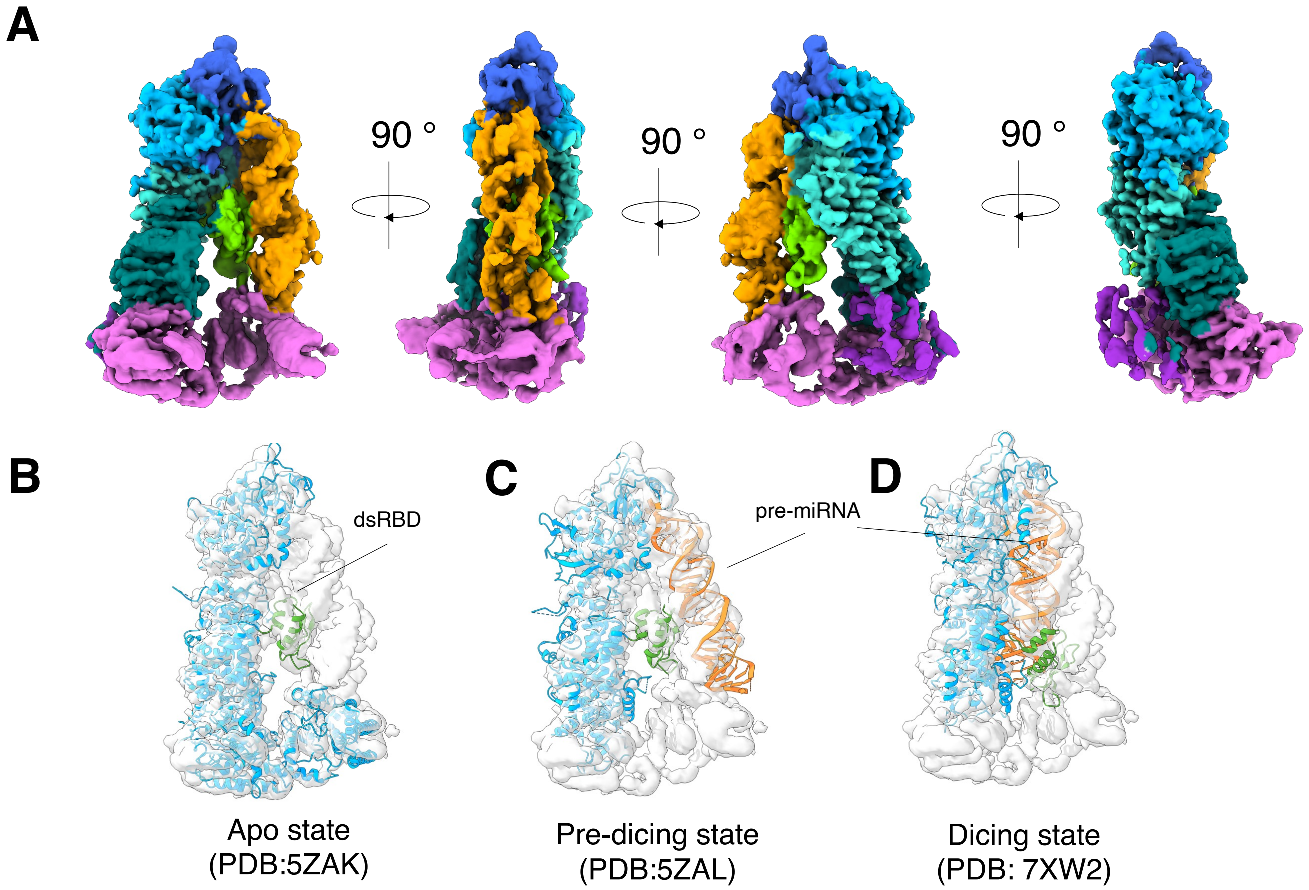


**Supplementary Figure S6.** (**A**) The composite map of Dicer-tRNA complex coloured by domain. (**B-D**) Different structural models are fitted into the cryo-EM map: (**B**) Dicer in apo-state (PDB: 5ZAK), (**C**) Dicer bound to pre-miRNA in a pre-dicing state, (**D**) and in a dicing state. Dicer is coloured in blue in (**B-D**) with the dsRBD highlighted in green and pre-miRNA in orange.

### tRNA-Ala +DMS

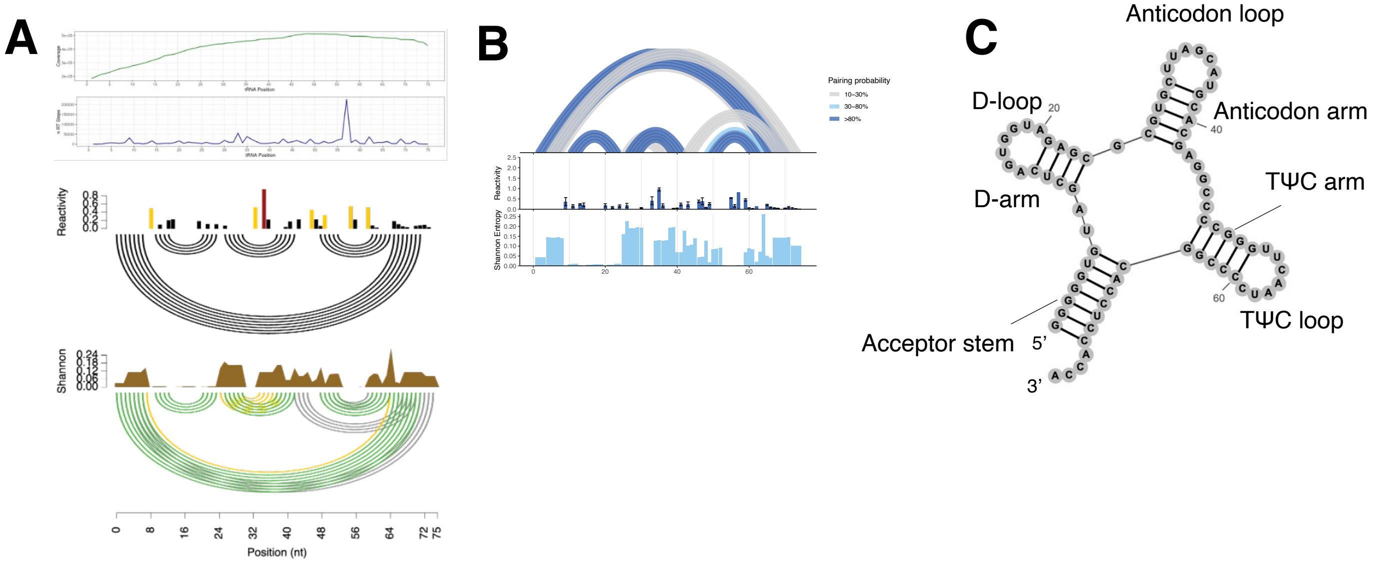


### tRNA-Ala +5NIA

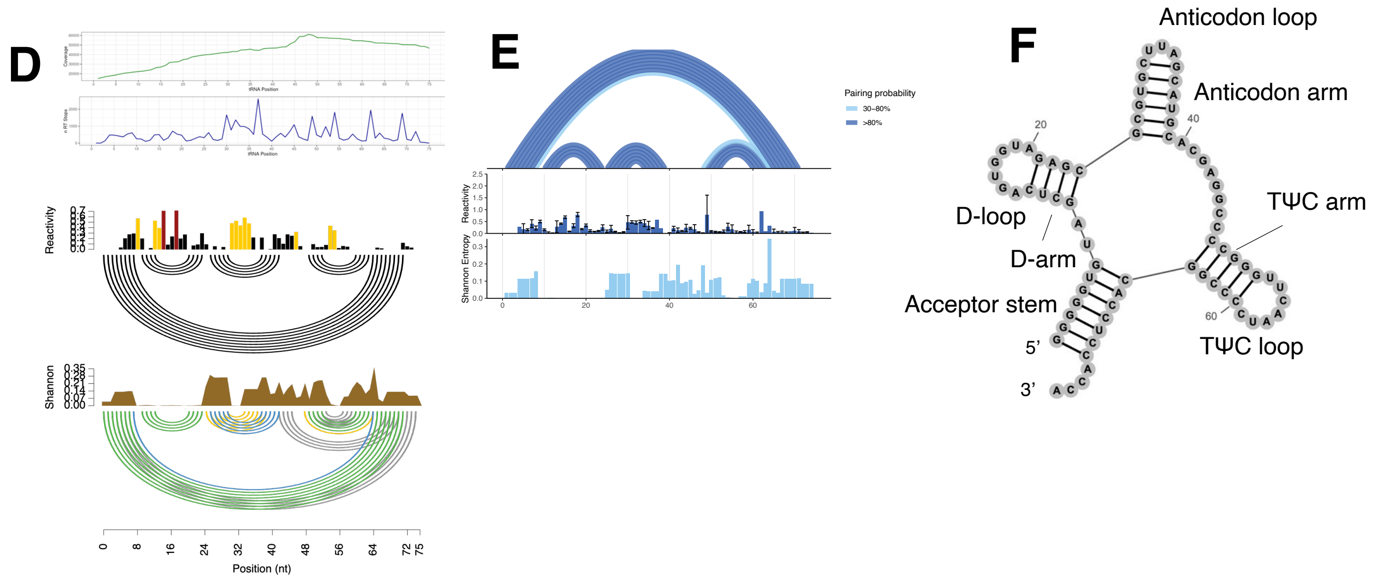


**Supplementary Figure S7.** The secondary structure of tRNA-Ala. Representative coverage, reverse transcription (RT) stops, reactivity and Shannon entropy plots of tRNA-Ala treated with (**A**) DMS or (**D**) 5NIA (*n=3)*. (**B**) Pairing probability, reactivity and Shannon entropy plot obtained by combined replicates of (**A**) and (**E**) for (**D**). Secondary structure of tRNA-Ala treated with DMS (**C**) or 5NIA (**F**), visualised with RNAcanvas.

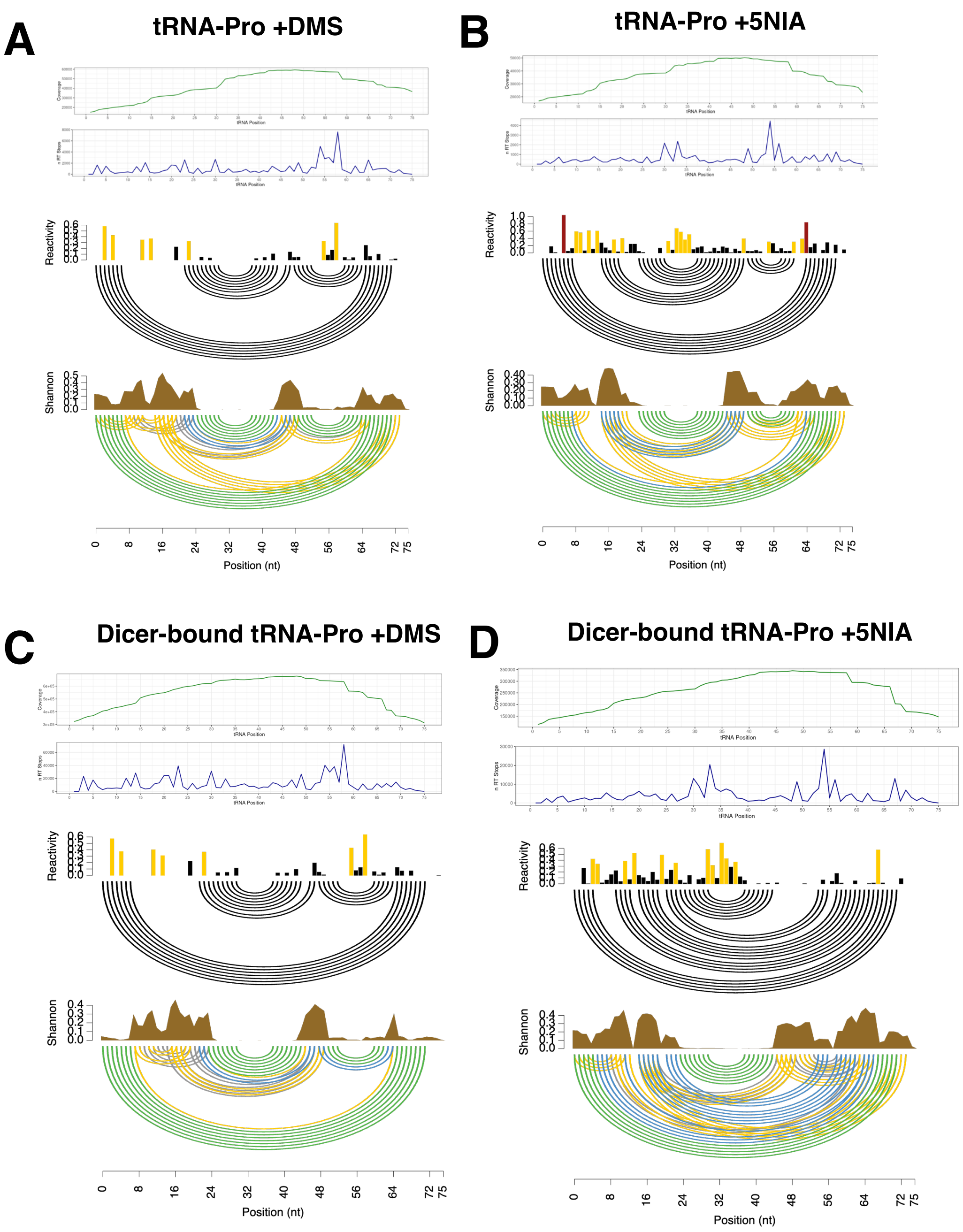


**Supplementary Figure S8.** The secondary structures of free and Dicer-bound tRNA-Pro. Representative coverage, reverse transcription (RT) stops, reactivity and Shannon entropy plots of free tRNA-Pro treated with (**A**) DMS or (**B**) 5NIA, and Dicer-bound tRNA-Pro treated with (**C**) DMS or (**D**) 5NIA (*n=3).*

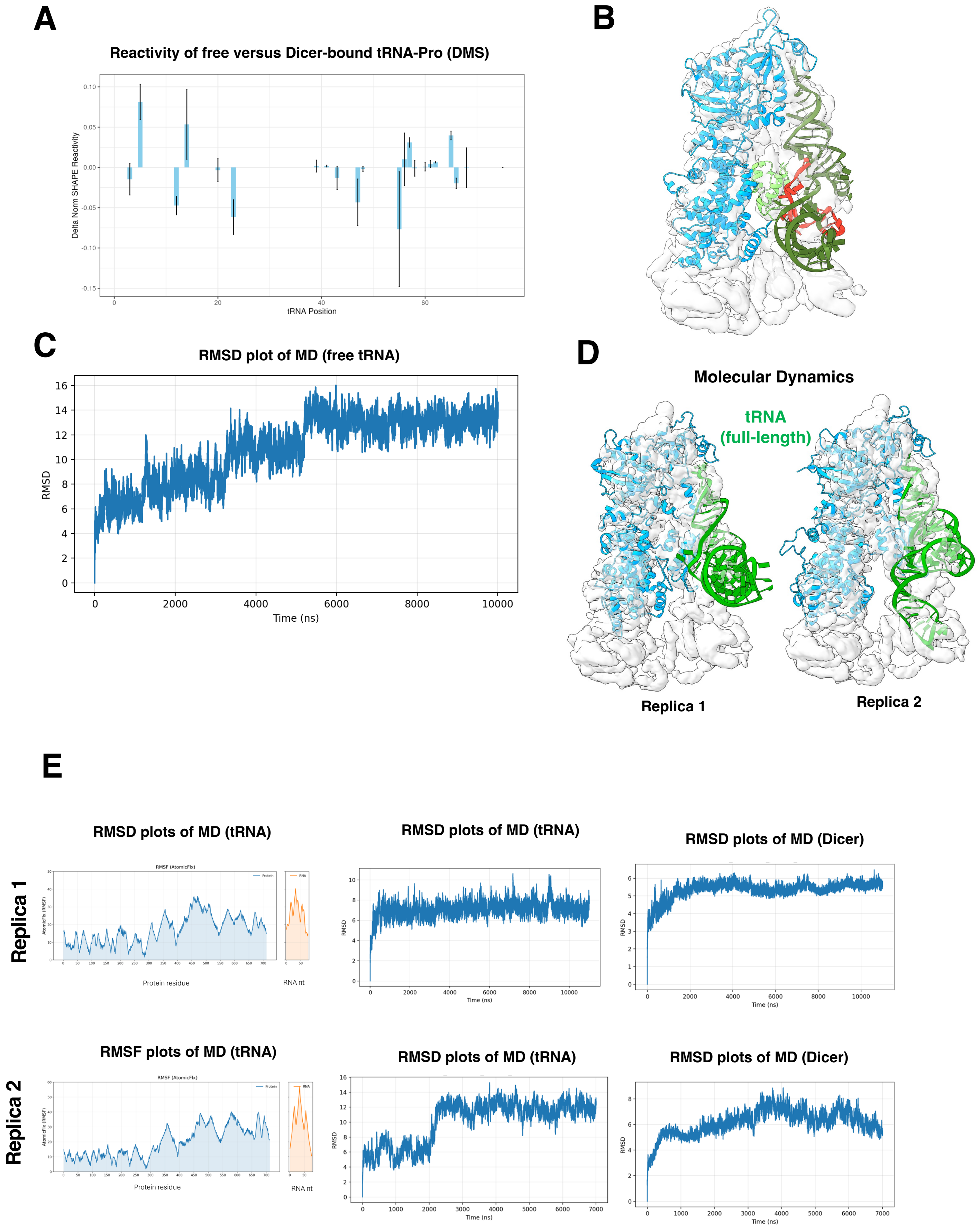


**Supplementary Figure S9.** (**A**) Changes in reactivity of free versus Dicer-bound tRNA-Pro treated with DMS. (**B**) Model of tRNA-Pro bound by Dicer. 3D tRNA model was obtained with trRosettaRNA guided by SHAPE of Dicer-bound tRNA (DMS); in red nucleotides with changes in SHAPE reactivity. (**C**) Root mean square deviation (RMSD) plot of MD of free tRNA-Pro *(n=2)*. (**D**) MD of Dicer-tRNA complex placed in cryo-EM map showing full length tRNA *(n=2)*. (**E**) Root mean square fluctuation (RMSF) and deviation (RMSD) plots of MD of the replica 1 and 2 of Dicer-tRNA-Pro complex *(n=2)*.

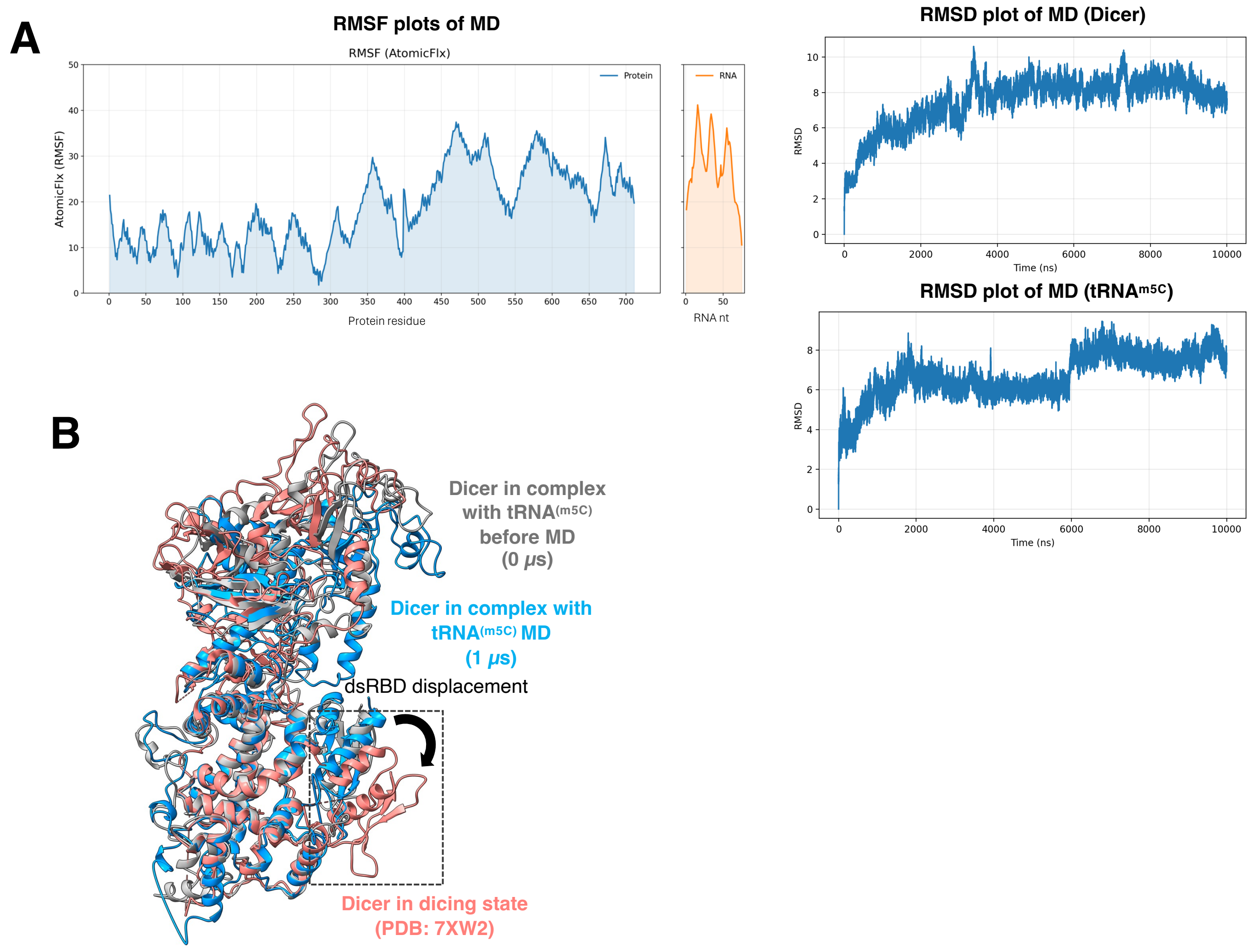


**Supplementary Figure S10.** (**A**) Representative root mean square fluctuation (RMSF) and root mean square deviation (RMSD) plots of molecular dynamics of Dicer-tRNA$^{(m5C)}$ *(n=2)*. (**B**) Structural comparison of structure of Dicer in dicing state (PDB: 7XW2) (pink) and Dicer in complex with tRNA$^{(m5C)}$ at 0 μs (grey) and 1 μs (blue) of molecular dynamics simulation.

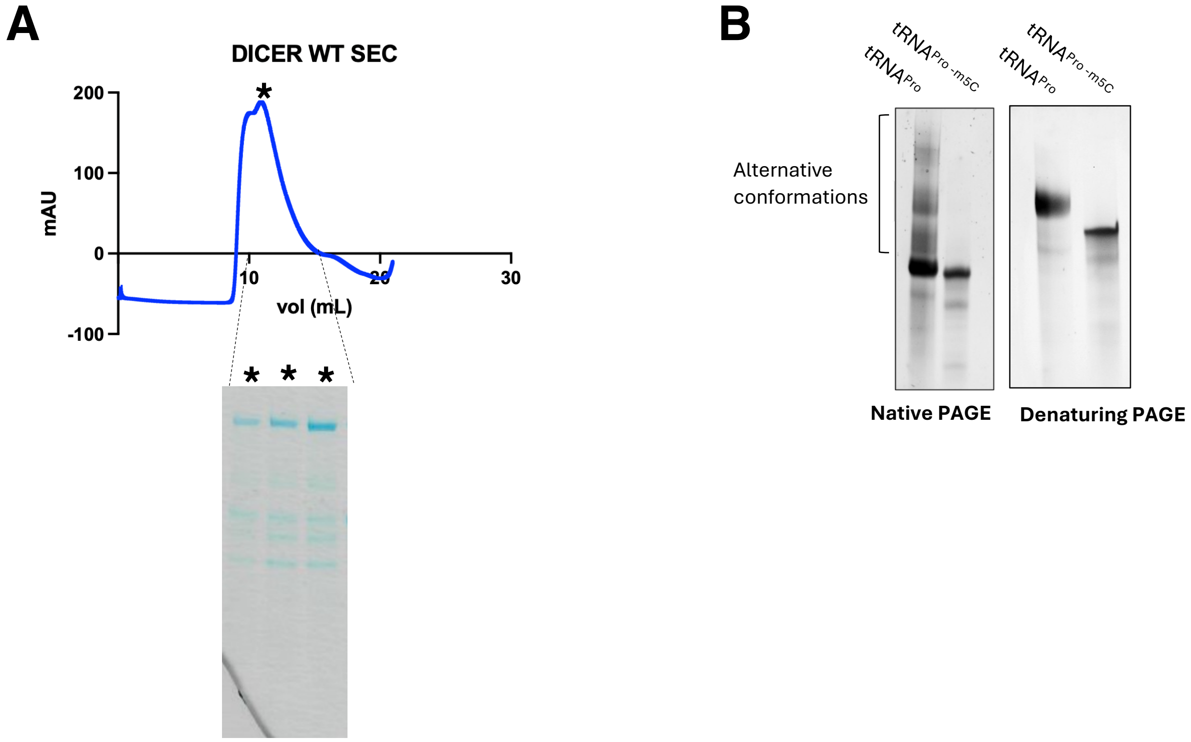


**Supplementary Figure S11.** (**A**) Size exclusion chromatography profile and reducing SDS-PAGE gel of purified Dicer wild type. (**B**) Native PAGE and denaturing PAGE of tRNA-Pro and tRNA-Pro$^{m5C}$.

| Name | Type | Sequence 5'3' | Experiment |
| --- | --- | --- | --- |
| Pro_3' | Northern Blot probe | TGGGGGCTCGTCCGGGATTTGAAC | Northern blots of Dicer cleavage assay |
| R1_Pro | Primer | TGGGGGCTCGTCCGG | *In vivo* Dicer RIP, Cleavage assay qRT-PCR |
| F1_Pro | Primer | GGCTCGTTGGTCTAGGGGTATG | *In vivo* Dicer RIP, Cleavage assay qRT-PCR |
| F1_SPINT1 | Primer | TAGCCTGTCTGTCTGCTAGG | Cleavage assay qRT-PCR |
| R1_SPINT1 | Primer | GATATTGCCCAACTACCCTCC | Cleavage assay qRT-PCR |
| F1_ALA | Primer | GGGGGTGTAGCTCAGTGGTAGAG | *In vivo* Dicer RIP, Cleavage assay qRT-PCR |
| R1_ALA | Primer | TGGTGGAGGTGCCGG | *In vivo* Dicer RIP, Cleavage assay qRT-PCR |
| F1_prelet7 | Primer | TGGGATGAGGTAGTAGGTTG | *In vivo* Dicer RIP qRT-PCR |
| R1_prelet7 | Primer | TAGGAAAGACAGTAGATTGTATAG | *In vivo* Dicer RIP qRT-PCR |
| ERCC00002_FWD_SPIN | Primer | GCTCACAGTATACGGGCGTC | *In vivo* Dicer RIP qRT-PCR |
| ERCC00002_REV_SPIN | Primer | ACCGTACAGCTCTGGAACCC | *In vivo* Dicer RIP qRT-PCR |
| RT primer SHAPE | Primer | CTACACGACGCTCTTCCGATCTNNNNNNNNNNTTTTTTTTTTTTTTTTTTTTTGG | SHAPE |
| | | | *In vitro* RNA transcription |
| T7-Pro TGG 3-5 | T7 DNA template | TAATACGACTCACTATAGGGGGCTCGTTGGTCTAGGGGTATGATTCTCGCTTTGGGTGCGAGAGGTCCCGGGTTCAAATCCCGGACGAGCCCCCA | *In vitro* RNA transcription |
| T7-Ala AGC 2-1 | T7 DNA template | TAATACGACTCACTATAGGGGGGGGTGTAGCTCAGTGGTAGAGCGCGTGCTTAGCATGCACGAGGCCCCGGGTTCAATCCCCGGCACCTCCACCA | *In vitro* RNA transcription |
| T7-Prelet7a | T7 DNA template | TAATACGACTCACTATAGGGTGGGATGAGGTAGTAGGTTGTATAGTTTTAGGGTCACACCCACCACTGGGAGATAACTATACAATCTACTGTCTTTCCTA | *In vitro* RNA transcription |

### Suppl. Table 1. List of oligos used in the paper

| Name | Manufacterer | Code | Experiment |
|---|---|---|---|
| Dicer | Proteintech | 20567 | PLA |
| NSUN2 | Proteintech | 66580 | PLA, WB |
| Dicer | Abcam | ab14601 | *In vitro* Dicer RIP and WB |

**Suppl. Table 2. List of antibodies used in the paper**
